# Single-cell multi-omics identify OTP and NOVA1 as regulators of leptin action in hypothalamic feeding circuits

**DOI:** 10.64898/2026.11.04.764326

**Authors:** Han L. Tan, Yuqi Tan, Jenelys Ruiz-Ortiz, Matt Paul, Zhaoyu Han, Leah Kelly, Anduena Toci, Kaja Plucinska, Yoko Tajima, Rani Shiao, Bora Lee, Anoj Ilanges, Robert B. Darnell, Paul Cohen, Jeffrey M. Friedman

**Affiliations:** Laboratory of Molecular Genetics, Howard Hughes Medical Institute, The Rockefeller University, New York, NY, USA; Department of Anatomy and Neurobiology, University of California Irvine School of Medicine, Irvine, CA, USA; Department of Microbiology and Immunology, Stanford University School of Medicine, Stanford, CA, USA; Bioinformatics Resource Center, The Rockefeller University, New York, NY, USA; Laboratory of Molecular Metabolism, The Rockefeller University, New York, NY, USA; Laboratory of Molecular Neuro-Oncology, Howard Hughes Medical Institute, The Rockefeller University, New York, NY, USA; Department of Biochemistry, Chonnam National University Medical School, Hwasun, Republic of Korea; Janelia Research Campus, Ashburn, VA, USA

## Abstract

Leptin acts directly and indirectly on multiple neuronal populations to regulate feeding and numerous other physiologic functions. However, our understanding of how a single circulating signal coordinates diverse neuronal populations across multiple circuits is limited. To further elucidate how leptin reshapes molecular states in its direct and indirect neural targets, we combined time-resolved single-nucleus transcriptomics and chromatin accessibility profiling to map leptin-induced transcriptional and chromatin state changes across individual neurons in two key hypothalamic nuclei: the arcuate nucleus (ARC), which contains the majority of leptin receptor (LepR) neurons, and the paraventricular hypothalamic nucleus (PVH), an established downstream target of ARC LepR neurons. We found that leptin treatment of leptin-deficient *ob/ob* mice does not alter neuronal identity, but instead induces coordinated, cell-type-specific transcriptional and regulatory state changes that propagate hierarchically from the ARC to downstream PVH neurons. Integrating RNA velocity, cis-regulatory analysis, and cell-cell communication inference revealed that leptin modulates hypothalamic circuit architecture by selectively strengthening specific interactions while pruning broadly distributed, low-specificity signaling. Although leptin broadly activated STAT3 signaling across LepR neuronal subtypes, integrated multi-omic analyses revealed that activation of STAT3 engages distinct downstream regulatory programs in different neuronal populations, often with opposing physiological functions. Functional analyses identified the homeobox transcription factor OTP, a leptin-responsive regulator selectively induced in AGRP neurons, as a critical mediator of leptin-mediated inhibition of hunger-promoting AGRP neuronal activity and feeding. In parallel, unbiased analysis of PVH neuronal state transitions shows that NOVA1, an RNA-binding protein primarily expressed in inhibitory neurons, is regulated by leptin and alterations of NOVA1 expression bidirectionally control food intake and body weight. Together, these findings define a multi-layered regulatory framework through which leptin regulates hypothalamic feeding circuits to control energy balance.

## Main

Leptin is the afferent signal in a negative feedback loop that regulates energy balance and numerous other physiological functions^1,2^. Leptin acts through the long isoform of the leptin receptor (LepRb) to activate canonical Janus kinase (JAK)-signal transducer and activator of transcription (STAT) pathways, particularly STAT3, in multiple hypothalamic neuronal populations^2,3^. However, leptin signaling suppresses the activity of hunger-promoting agouti-related peptide (AGRP)/LepRb neurons while activating satiety-promoting proopiomelanocortin (POMC)/LepRb and basonuclin 2 (BNC2)/LepRb neurons. Thus, leptin elicits different effects (i.e., neural activation vs. inhibition) in these populations despite the fact that all three express the same receptor and engage the canonical LepR-STAT3 signaling pathway. This raises the fundamental question of how a shared LepR-STAT3 signaling pathway generates distinct, and in some cases opposing, cellular responses^3–5^. In addition, how leptin-responsive signals propagate to downstream non-LepR neurons and how these distributed neuronal populations are organized into specific circuits over time to shape feeding behavior has not been systematically evaluated. A fuller understanding of these questions is of basic importance and has therapeutic implications because the glucagon-like peptide 1 receptor (GLP1R) agonists require intact leptin signaling, as these drugs are ineffective in leptin-deficient mice, leptin receptor-deficient mice, and in mice lacking LepR in GLP1R-expressing neurons^6–9^. This suggests that leptin signaling may induce neuronal state changes necessary for the response to satiety-promoting cues, highlighting the importance of understanding how leptin organizes feeding circuits at the molecular and cellular levels.

Leptin acts on first-order LepR-expressing neurons in the arcuate nucleus (ARC) of the hypothalamus, a key site of leptin action, which relay metabolic state information to downstream feeding circuits^2^. The paraventricular hypothalamic nucleus (PVH) is a major downstream target of ARC LepR neurons and serves as a critical output node controlling feeding behavior, energy expenditure, autonomic function, and neuroendocrine responses through its projections to the hindbrain, spinal cord, and neuroendocrine centers^10,11^. To define how leptin signaling generates distinct neuronal responses and propagates across hypothalamic feeding circuits, we performed time-resolved single-nucleus RNA sequencing (snRNA-seq) and chromatin accessibility profiling (snATAC-seq) of ARC and PVH neurons following leptin treatment.

Here we show that leptin induces coordinated, cell-type-specific transcriptional and regulatory state transitions that propagate hierarchically from ARC LepR neurons to downstream hypothalamic populations. Integrative analyses further revealed that LepR-STAT3 signaling engages distinct downstream transcriptional programs in different LepR neuron subtypes. These analyses identified many leptin-regulated genes, including the transcription factor orthopedia homeobox (OTP), which is necessary for leptin-dependent inhibition of hunger-promoting AGRP/LepR neurons. In contrast, LepR-STAT3 signaling regulated different sets of downstream genes in satiety-promoting BNC2/LepR and POMC/LepR neurons. We also found that leptin induced substantial state transitions in downstream hypothalamic populations not expressing LepR in the PVH, with changes of even greater magnitude than were observed in ARC neurons, suggesting a hierarchical circuit reorganization. Unbiased analyses identified neuro-oncological ventral antigen 1 (NOVA1), an RNA-binding protein previously studied primarily in inhibitory neurons^12,13^, as a regulator of leptin-responsive state transitions in PVH neurons. Changes in NOVA1 expression in PVH bidirectionally control feeding and body weight. Together, these findings reveal novel cellular and molecular mechanisms in both primary and downstream target neurons and establish a molecular and cellular framework for understanding how leptin organizes hierarchical, cell-type-specific state transitions across distributed hypothalamic circuits to coordinate energy balance.

## Results

### A multi-omic atlas of leptin-responsive ARC/PVH neurons

To define the cellular and molecular basis of leptin action across hypothalamic feeding circuits, we performed multiplexed snRNA-seq and snATAC-seq on flow cytometry-enriched neuronal nuclei from microdissected ARC and PVH of adult wild-type (WT) controls and age-matched leptin-deficient *ob/ob* mice at baseline and after leptin treatment for 3, 27, and 75 hours ^14,15^ (**Fig. 1a**). For the 27-hour and 75-hour time points, leptin was administered every 8 hours to maintain sustained and consistent leptin exposure. In each case, tissues were collected 3 hours after the final injection to allow comparison across matched time windows post-leptin treatment. This experimental paradigm enabled us to map leptin’s effects over a range of circulating levels and identify synchronized activation of leptin-responsive pathways.

**Fig. 1:**
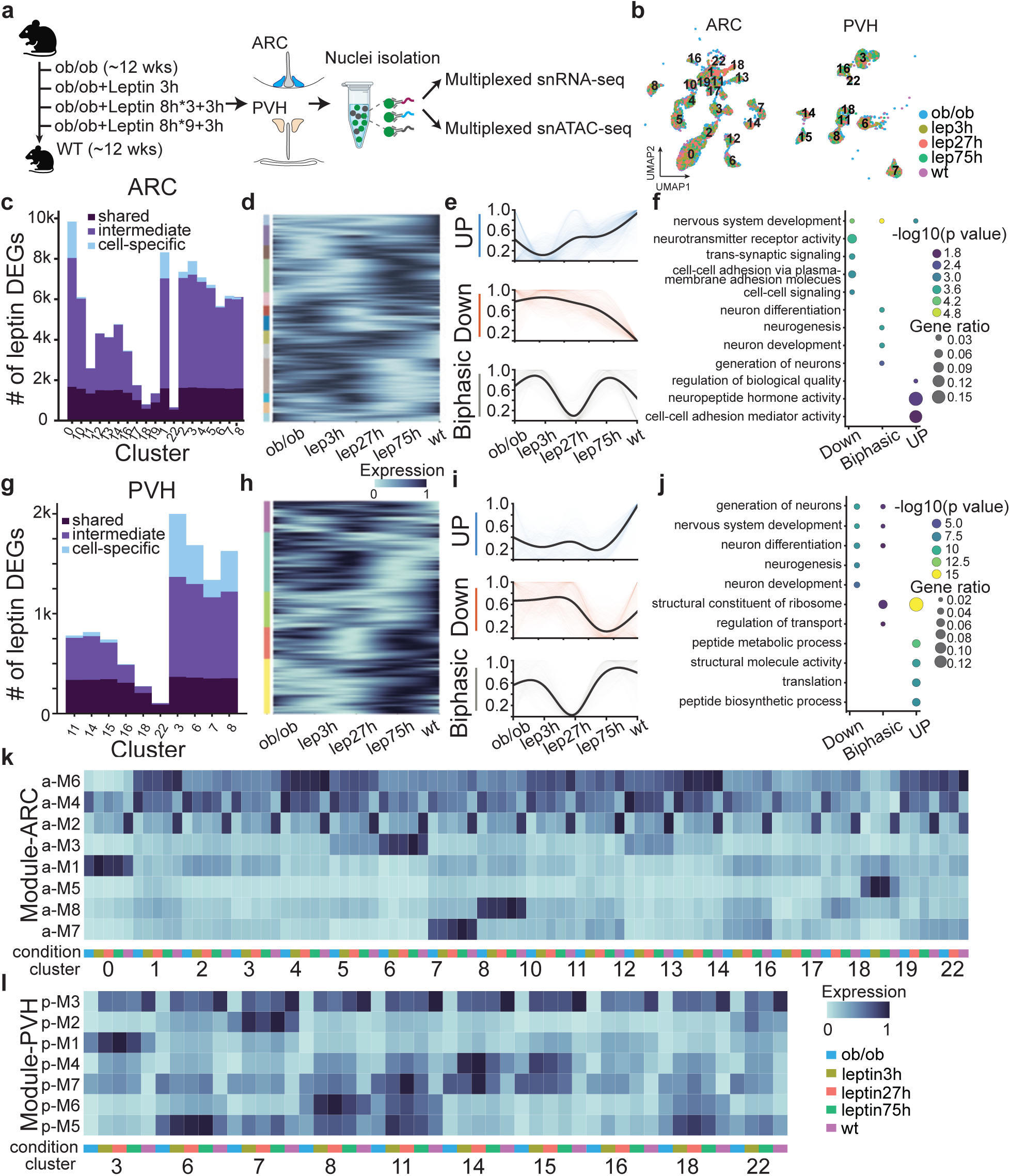
A multi-omic atlas of leptin-responsive ARC/PVH neurons. **a**, Experimental design and workflow for multiplexed snRNA-seq and snATAC-seq profiling of ARC and PVH neurons from ob/ob, leptin-treated ob/ob, and WT mice. **b**, UMAP visualization of ARC and PVH neuronal populations colored by experimental condition; numbers indicate neuronal clusters. **c,g,** Number of leptin-responsive differentially expressed genes (lepDEGs) across ARC (**c**) and PVH (**g**) neuronal clusters. **d,h,** Heatmaps showing expression patterns of lepDEGs in ARC (**d**) and PVH (**h**) neurons across experimental conditions. Genes are grouped according to their temporal expression dynamics (left colored bars). **e,i,** Examples of lepDEG clusters from **d** and **h**. Colored lines indicate expression trajectories of individual lepDEGs, and black lines indicate the mean expression trajectory of all genes within each cluster. **f,j,** Gene Ontology (GO) enrichment analysis of the lepDEG clusters shown in **e** and **i. k,l,** Mean scaled expression of genes within each module across ARC (**k**) and PVH (**l**) neuronal clusters and experimental conditions. Columns represent neuronal clusters and conditions.

After quality control, we obtained 24,198 nuclei for snRNA-seq and 33,899 nuclei for snATAC-seq. The vast majority of nuclei were neuronal, based on enrichment of neuronal marker genes and depletion of non-neuronal signatures (**Extended Data Fig. 1**). Unsupervised Leiden clustering of the snRNA-seq datasets identified 21 neuronal clusters in ARC of which 19 were matched those reported elsewhere (**Fig. 1b**). This classification recovered several well established leptin responsive neuronal populations, including AGRP/LepR, POMC/LepR and BNC2/LepR neurons^16,17^, and also identified previously uncharacterized populations such as lymphoid enhancer-binding factor 1 (LEF1) neurons (**Extended Data Fig. 2a**).

We also identified 23 neuronal clusters in PVH, 10 of which were molecularly defined excitatory PVH populations, and all of which expressed the canonical PVH marker *Sim1* (**Extended Data Fig. 2b**). While GABAergic neurons were also detected in PVH, we did not find any PVH-specific marker genes in these populations, and these clusters expressed markers broadly shared with adjacent hypothalamic regions (**Extended Data Fig. 2c,d**). We thus focused subsequent analyses of PVH on the excitatory populations expressing *Sim1*. This classification recovered established neuronal populations, including BDNF and GLP1R neurons, and also identified a previously uncharacterized neuronal population expressing the piezo-type mechanosensitive ion channel component 2 (PIEZO2) (**Extended Data Fig. 2b**).

In ARC and PVH, neuronal identities and cluster organization remained unchanged across all experimental conditions, and we did not observe any differences between WT and *ob/ob* mice, nor were there major treatment-specific clusters following leptin administration (**Fig. 1b**). These findings indicate that leptin primarily modulates neuronal state rather than fundamentally altering cell-type identity.

To characterize leptin-induced transcriptional responses, we performed differential expression analysis across the leptin treatment time course for each neuronal cluster and identified thousands of leptin-responsive genes (LepDEGs). These differentially expressed genes included *Fos* (*cFos*), which, consistent with prior data, was increased in POMC/LepR and reduced in AGRP/LepR neurons after leptin treatment. We also observed changes in several components of the leptin signal transduction pathway, including *Lepr*, *Stat3*, and *Socs3*, with notable differences among them ^2^ (**Extended Data Fig. 3a-c**). The induction of *Stat3* and its downstream target *Socs3* in AGRP/LepR, POMC/LepR, and BNC2/LepR neurons was consistent with activation of canonical leptin signaling, whereas *Lepr* expression exhibited cell-type-specific changes. More broadly, LepDEGs showed diverse expression patterns, with some genes being broadly regulated across multiple neuronal populations and others being highly restricted to specific neuronal types (**Fig. 1c,g**), revealing both shared and population-specific transcriptional responses to leptin across hypothalamic circuits. To further define these spatial and temporal patterns of leptin- responsive gene expression, we grouped LepDEGs according to their expression dynamics across the leptin treatment time course. This analysis identified multiple response patterns, including progressively induced, progressively repressed, and biphasic gene programs ( **Fig. 1d,e,h,i**). Gene ontology analysis linked these temporal programs to several distinct biological pathways, including neuronal development, trans-synaptic signaling, neuropeptide hormone activity, and peptide biosynthetic processes, with differing contributions across ARC and PVH neurons (**Fig. 1f,j**).

Higher-order transcriptional programs underlying leptin-responsive gene expression beyond individual DEGs were identified using Hotspot analysis to derive co-expression modules across ARC and PVH neurons^18^ (**Extended Data Fig. 4a-d**). These modules grouped coordinately expressed genes and captured both stable neuronal identity programs and dynamic leptin-responsive transcriptional states^18^. As above, some modules were broadly regulated across multiple neuronal populations, whereas others exhibited strong cell-type specificity or minimal leptin responsiveness (**Fig. 1k,l, Extended Data Fig. 4a,c**). For example, ARC Module 1 (a-M1, AGRP/LepR identity module) was strongly enriched in AGRP/LepR neurons which expressed canonical AGRP/LepR-associated genes including *Agrp*, *Npy*, *Otp*, and *Acvr1c*, representing a core AGRP/LepR neuronal identity program that was largely preserved across conditions with only modest leptin-dependent modulation (**Fig. 1k, Extended Data Fig. 4a**). (Note, the function of *Otp* in AGRP/LepR neurons is described in detail below). In contrast, ARC-wide Module 2 (a-M2), which increased following leptin treatment across most ARC neuronal populations, was enriched for genes associated with mitochondrial energy metabolism, whereas Module 4 (a-M4), which decreased, was enriched for genes associated with neuronal signaling and development (**Fig. 1k, Extended Data Fig. 4a,b**). The response to leptin in PVH was similar, where certain co-expression modules were restricted to specific neuronal populations, while others were broadly expressed across multiple PVH clusters (**Fig. 1l, Extended Data Fig. 4c,d**). Together, these analyses reveal that leptin engages both stable neuronal identity programs and activates dynamic cell-type-specific transcriptional programs across many hypothalamic populations. Specific examples of each of these patterns are described in greater detail below.

### Cell-type-specific neuronal state transitions induced by leptin

We further refined the analysis of leptin-responsive transcriptional programs across ARC and PVH neurons (**Fig. 1**) using RNA velocity analysis, which infers the direction of transcriptional state changes from RNA splicing dynamics, to examine how leptin reshapes neuronal states across time and space^19^. To quantify these transitions across neuronal populations, we summarized velocity-derived pseudotime values for each ARC and PVH cluster (**Fig. 2a,b**). Across most neuronal populations, with the exception of ARC clusters 16 and 18, cells from *ob/ob*, leptin-treated, and WT conditions showed a consistent ordered progression, with leptin-treated cells positioned between leptin-deficient and WT states (the characteristics of Clusters 16 and 18 are discussed further below). Differences among the three time points for leptin treatment (3 h, 27 h, and 75 h) were relatively modest compared to the greater separation between neurons from untreated *ob/ob* and WT mice, suggesting that leptin rapidly induces major transcriptional state changes that are stable over time. Leptin receptor was not expressed in PVH, and pseudotime shifts were generally larger in PVH neurons than in ARC neurons, indicating broader transcriptional remodeling among PVH populations. This is consistent with a hierarchical circuit organization in which leptin-responsive signals propagate from ARC to downstream PVH neurons.

**Fig. 2:**
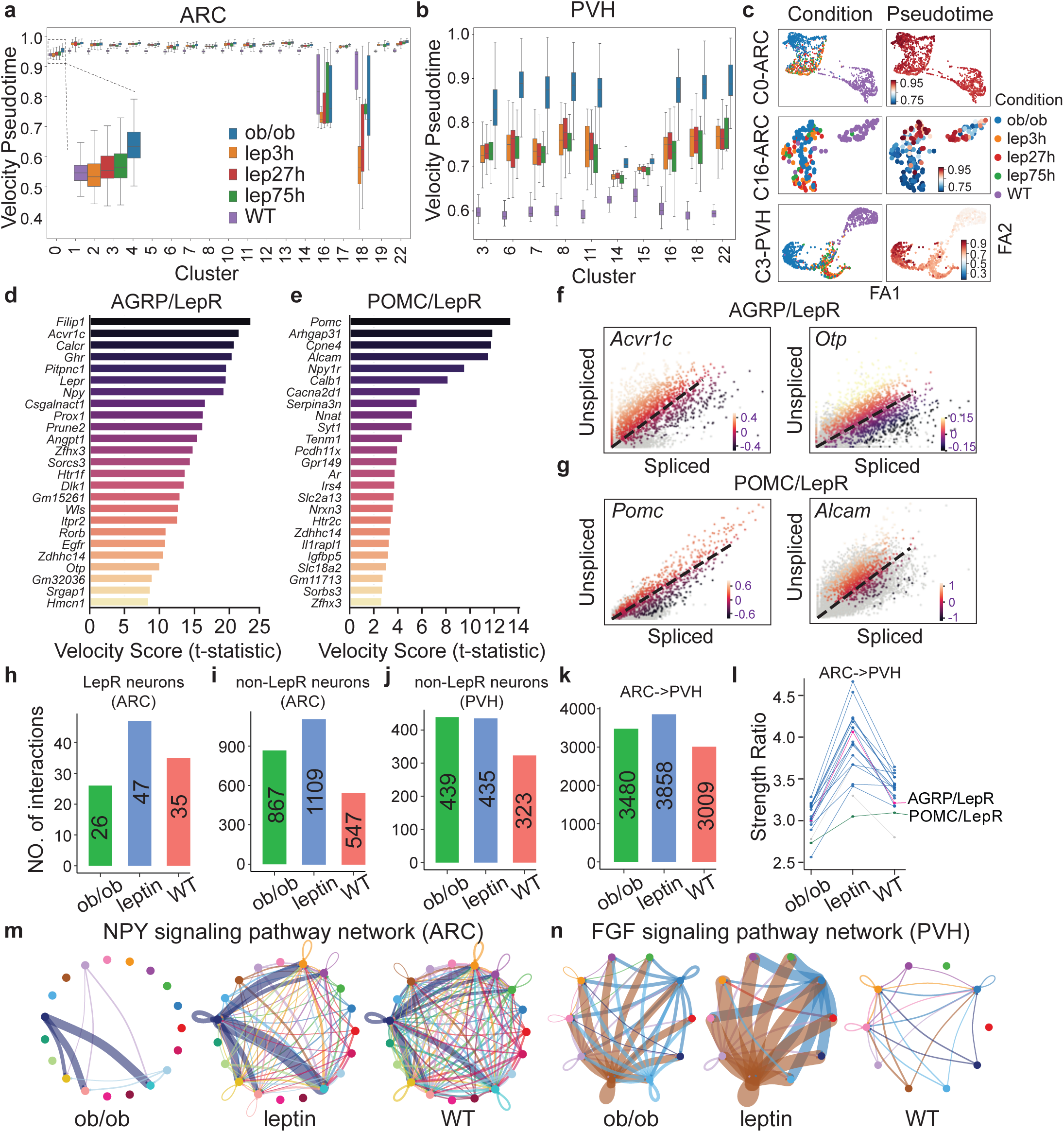
Cell-type-specific neuronal state transitions induced by leptin. **a,b,** RNA velocity pseudotime distributions across ARC (**a**) and PVH (**b**) neuronal clusters under ob/ob, leptin-treated ob/ob, and WT conditions. Boxes indicate median and interquartile range. **c**, Force-directed graph projections of representative neuronal populations (ARC clusters 0/16 and PVH cluster 3) colored by experimental condition (left) or velocity pseudotime (right). **d,e,** Top velocity-associated genes ranked by velocity score in AGRP/LepR neurons (**d**) and POMC/LepR neurons (**e**). **f,g,** Phase portraits of representative velocity-associated genes in AGRP/LepR neurons (**f**) and POMC/LepR neurons (**g**). **h-j,** Number of inferred cell-cell interactions among LepR neurons (**h**) and non-LepR neurons (**i**) in the ARC and PVH neurons (**j**) across conditions. **k,** Total number of inferred ARC-to-PVH interactions across conditions. **l,** Interaction strength ratios for inferred ARC-to-PVH signaling pathways across conditions. **m,n,** Representative inferred signaling networks for the NPY pathway in ARC neurons (**m**) and the FGF pathway in PVH neurons (**n**) under ob/ob, leptin-treated, and WT conditions. Node colors indicate neuronal clusters; edge width corresponds to inferred interaction strength.

To visualize these transitions at single-cell resolution, we generated force-directed graph projections of representative neuronal populations (**Fig. 2c**). Leptin-responsive populations, including ARC AGRP/LepR (C0) neurons and PVH corticotropin-releasing hormone (CRH) neurons (C3), recapitulated the ordered progression observed in the global analysis, with leptin-treated cells occupying intermediate positions between *ob/ob* and WT states. In contrast, non-responsive populations such as ARC cluster 16 (C16) lacked coherent directional organization and exhibited extensive intermixing of cells across all conditions (**Fig. 2c**), indicating minimal leptin-dependent remodeling.

We next ranked genes within each neuronal population based on their contribution to RNA velocity. Velocity-associated genes represent genes whose spliced and unspliced transcript dynamics most strongly contribute to the inferred future transcriptional states of individual cells. These genes were highly cell-type specific and revealed distinct molecular programs across ARC and PVH neurons (**Supplementary Tables 1,2**). In AGRP/LepR neurons, the top velocity-associated genes included neuropeptide (*Npy*), receptor genes (*Lepr*, *Ghr*, *Calcr*, and *Acvr1c*), and the transcription factor *Otp* (**Fig. 2d**), highlighting distinct classes of candidate molecular regulators underlying leptin-responsive state transitions. In contrast, POMC/LepR neurons exhibited a different set of velocity-associated genes, including the neuropeptide *Pomc*, calcium-associated genes (*Calb1* and *Cacna2d1*), the serotonin receptor (*Htr2c)*, and the insulin signaling adaptor protein (*Irs4)* (**Fig. 2e**), demonstrating that leptin engages a distinct molecular program in POMC/LepR neurons compared with the AGRP/LepR population. Phase portrait analyses further supported these transitions at the gene level. In AGRP/LepR neurons, *Acvr1c* and *Otp* displayed positive velocity signatures consistent with ongoing transcriptional regulation, further suggesting that these genes are actively upregulated during leptin-responsive state transitions (**Fig. 2f**). In contrast, POMC/LepR neurons exhibited induction of genes including *Pomc* and *Alcam* along a distinct trajectory (**Fig. 2g**), further supporting divergent molecular programs across LepR neuronal subtypes. Together, these analyses define cell-type-specific molecular trajectories underlying leptin-responsive neuronal remodeling across ARC and PVH populations.

To examine how these neuronal state changes are coordinated at the circuit level, we applied CellChat to infer ligand-receptor-mediated intercellular communication networks within and between ARC and PVH neurons^20^. Because transcriptional differences at each of the three time points of leptin treatment (3h, 27h, and 75h) were modest relative to the separation between leptin-deficient *ob/ob* mice and WT animals, data from all three time points were pooled for downstream interaction analyses. This analysis recovered known hypothalamic interactions, including BNC2/LepR→AGRP/LepR and AGRP/LepR→POMC/LepR signaling^3,21^, and identified additional novel candidate interactions such as BNC2/LepR→growth hormone-releasing hormone (GHRH) neurons (**Extended Data Fig. 5**). Given the reported orexigenic function of GHRH neurons^22^ and established links between leptin, melanocortin-3 receptor (MC3R) signaling, and the GHRH-GH axis^23^, this interaction may represent an additional pathway through which satiety-promoting BNC2/LepR neurons influence feeding behavior and growth. Quantitative analysis revealed marked cell-type-specific changes in inferred communication following leptin treatment. LepR-expressing AGRP (cluster 0), POMC (cluster 3), and BNC2 (cluster 17) neurons exhibited the lowest numbers of inferred interactions in *ob/ob* mice but showed a pronounced increase following leptin treatment (**Fig. 2h; Extended Data Fig. 5**). In contrast, non-LepR ARC neurons and PVH neurons exhibited the opposite pattern, with the highest interaction values in *ob/ob* mice and a reduction following leptin treatment (**Fig. 2i,j; Extended Data Fig. 5**). Analysis of inter-regional ARC-to-PVH communication revealed a similar pattern. ARC-to-PVH interaction numbers were elevated in *ob/ob* mice and decreased following leptin treatment (**Fig. 2k; Extended Data Fig. 5**). However, interaction specificity ratios increased after leptin treatment across most ARC neuronal populations, indicating a shift from broadly distributed communication patterns in *ob/ob* mice toward a smaller set of more selective ARC-to-PVH signaling interactions after leptin treatment (**Fig. 2l**). At the pathway level, leptin induced region-specific shifts in dominant signaling programs. In the ARC, neuropeptide Y (NPY) signaling was minimal in *ob/ob* mice and progressively increased following leptin treatment toward WT levels (**Fig. 2m**). In contrast, fibroblast growth factor (FGF) signaling in the PVH was highest in *ob/ob* mice and markedly lower in WT animals (**Fig. 2n**). Together, these findings suggest that leptin reorganizes hypothalamic communication networks and engages distinct signaling pathways from ARC to PVH neurons.

### Shared LepR-STAT3 signaling engages distinct regulatory programs

Having identified distinct leptin-responsive transcriptional programs and neuronal state transitions across hypothalamic populations, we next sought to define the regulatory mechanisms underlying these cell-type-specific responses. To link transcriptional changes with upstream regulatory DNA accessibility and sequence, we integrated snRNA-seq and snATAC-seq datasets from ARC and PVH neurons. Clustering of snATAC-seq profiles based on chromatin accessibility, followed by annotation using gene activity scores and marker genes derived from snRNA-seq, identified corresponding neuronal populations across modalities. Gene activity inferred from chromatin accessibility showed strong concordance with transcriptomic cell identities (**Fig. 3a,b**), establishing a unified cellular framework for identifying leptin target genes and their regulatory mechanisms.

**Fig. 3:**
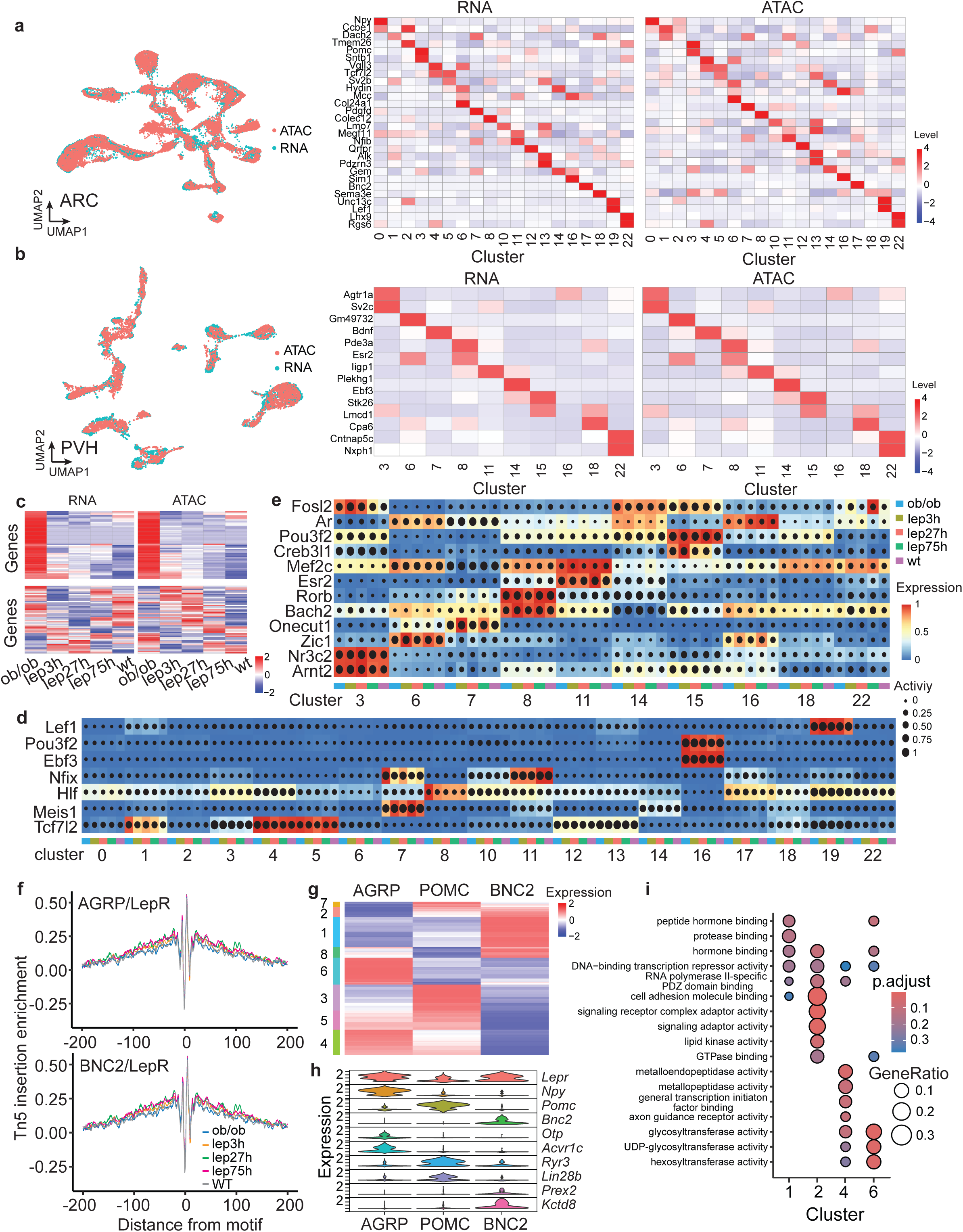
Shared LepR-STAT3 signaling engages distinct regulatory programs. **a,b,** Integration of snRNA-seq and snATAC-seq datasets from ARC (**a**) and PVH (**b**) neurons. Left, UMAP visualization of neuronal populations identified by snRNA-seq and snATAC-seq. Right, heatmaps show the expression of cluster marker genes in the snRNA-seq data and corresponding gene activity scores inferred from the snATAC-seq data. **c,** Representative genes showing concordant (top) and discordant (bottom) RNA and chromatin accessibility dynamics across the leptin treatment time course. **d,e,** SCENIC+-inferred regulon activity across ARC (**d**) and PVH (**e**) neuronal populations and experimental conditions. Dot color indicates scaled TF expression and dot size indicates regulon activity. **f,** Aggregate chromatin accessibility profiles centered on STAT3 motif sites in AGRP/LepR and BNC2/LepR neurons across experimental conditions. **g,** Heatmap showing expression of SCENIC+-predicted STAT3 target genes across AGRP/LepR, POMC/LepR, and BNC2/LepR neuronal populations. Genes are grouped into distinct expression modules based on shared expression patterns (left colored bars). **h,** Expression of representative STAT3 target genes across AGRP/LepR, POMC/LepR, and BNC2/LepR neurons, including genes enriched in AGRP/LepR (*Otp* and *Acvr1c*), POMC/LepR (*Ryr3* and *Lin28b*), and BNC2/LepR (*Prex2* and *Kctd8*) neurons. **i,** Gene Ontology molecular function enrichment analysis of the STAT3 target gene modules shown in **g**.

We next constructed a cis-regulatory map linking chromatin accessibility to gene expression in a cell-type-resolved manner. Each snATAC-seq nucleus was matched to its nearest neighbor in the snRNA-seq dataset, enabling identification of putative cis-regulatory elements (CREs) whose accessibility correlated with expression of nearby genes (<250 kb). This analysis generated a genome-wide set of CRE-gene associations across ARC and PVH neurons (**Extended Data Fig. 6a-d**). For many leptin-responsive genes, accessibility at linked cis-regulatory elements broadly tracked transcriptional dynamics across the leptin treatment time course (**Fig. 3c**), consistent with coordinated chromatin and transcriptional remodeling during leptin treatment. However, some genes exhibited transcriptional changes that were weakly correlated or discordant with accessibility dynamics (**Fig. 3c**), suggesting that leptin-responsive transcription can also occur independent of chromatin remodeling.

To identify transcriptional regulators underlying these coordinated chromatin and transcriptional programs, we applied SCENIC+ to the integrated multi-omic dataset^24^. This analysis links accessible regulatory elements with enriched transcription factor (TF) motifs and co-expressed target genes to infer cell-type-specific regulon activity. Many TF regulons were stably detected across experimental conditions, consistent with core identity programs that maintain neuronal subtype specification. Examples included *Lef1* in ARC cluster 19, *Ebf3* in ARC cluster 16, and *Nr3c2* in PVH cluster 3 (**Fig. 3d,e**). In contrast, a subset of TF regulons displayed dynamic leptin-dependent changes in activity. Notably, *Tcf7l2* regulon activity decreased in ARC cluster 10 following leptin treatment (**Fig. 3d**), implicating this diabetes-associated transcription factor in hypothalamic leptin responses. Additional TFs, including *Meis1* in ARC cluster 7 and *Ar* in PVH cluster 16, showed increased regulon activity following leptin treatment, revealing candidate leptin-responsive regulators operating in distinct neuronal populations (**Fig. 3d,e**).

Because leptin signaling converges on STAT3 activation across multiple LepR neuronal populations^2^, we next asked whether STAT3 signaling engages common or divergent downstream regulatory programs across distinct LepR neurons. STAT3 motif accessibility and regulon activity were significantly increased across all LepR neuronal subtypes following leptin treatment (**Fig. 3f**), consistent with canonical activation of LepR-STAT3 signaling. However, despite shared STAT3 activation, inferred STAT3 regulatory networks using SCENIC+ analysis diverged markedly across LepR neuronal populations (**Fig. 3g**; **Supplementary Tables 3**). In hunger-promoting AGRP/LepR neurons, predicted STAT3 targets included *Otp* and *Acvr1c*, which were selectively enriched in this population and minimally expressed in satiety-associated BNC2/LepR and POMC/LepR neurons (**Fig. 3g,h**). In POMC/LepR neurons, predicted STAT3 targets included *Ryr3* and *Lin28b*, which were enriched in this population relative to AGRP/LepR and BNC2/LepR neurons (**Fig. 3g,h**). In contrast, in BNC2/LepR neurons, STAT3 was predicted to regulate a distinct set of targets, including *Prex2* and *Kctd8*, which were weakly expressed or absent in AGRP/LepR neurons (**Fig. 3g,h**). Gene ontology analysis further revealed functional specialization of STAT3-associated regulatory programs across LepR subtypes. Predicted STAT3 target genes in AGRP/LepR neurons were associated with molecular functions related to glycosyltransferase activity, hexosyltransferase activity, and metalloendopeptidase activity, whereas predicted STAT3 targets in BNC2/LepR neurons were associated with hormone binding, protease binding, and cell-adhesion molecule binding (**Fig. 3i**). Together, these findings indicate that a shared leptin-STAT3 signaling pathway engages distinct downstream regulatory programs across heterogeneous LepR neuronal populations. We next used this information to evaluate the function of specific genes identified using these analyses.

### OTP is required for leptin-mediated inhibition of AGRP neurons

Our integrated transcriptomic, chromatin accessibility, and regulatory network analyses identified multiple candidate regulators of leptin action that were selectively engaged across distinct neuronal populations. To prioritize candidate regulators for functional validation, we focused on leptin-responsive genes that were consistently prioritized across multiple complementary analyses, including differential expression, RNA velocity, regulatory network inference, and, importantly, evidence of STAT3 regulation.

Among these candidates, the homeobox transcription factor OTP emerged as a prominent regulator in AGRP/LepR neurons. OTP was consistently prioritized across complementary analyses: it ranked highly among velocity-associated genes, was identified as a predicted STAT3 target, and exhibited selective expression in AGRP/LepR neurons relative to other LepR populations (**Fig. 2d**, **3i; Extended Data Fig. 6f**). Moreover, while the function of other genes with similar patterns of expression has been previously evaluated (for example, *Npy*, *Lepr*, *Ghr*, *Calcr*, and *Acvr1c)*, the role of *Otp* in leptin action has not. Together, these observations suggested that OTP may function as a downstream effector of leptin signaling in hunger-promoting AGRP/LepR neurons, prompting us to further test its function in AGRP/LepR neurons of WT mice. While OTP is best known for its developmental roles in hypothalamic cell specification^25,26^, as mentioned, its function in mature AGRP/LepR neurons in adults and in leptin signaling has not been evaluated.

Consistent with a possible role as a leptin-responsive regulator, leptin robustly increased *Otp* expression selectively in AGRP/LepR neurons (**Fig. 4a**). To determine whether OTP is functionally required for leptin action in AGRP/LepR neurons, we selectively disrupted *Otp* in adult AGRP neurons using a CRISPR-Cas9 approach. AGRP-Cre mice were crossed with LSL-Cas9-GFP mice, and an adeno-associated virus (AAV) encoding sgRNAs targeting *Otp* (sgOtp) or control sgRNAs (sgCtrl) was bilaterally injected into the ARC (**Extended Data Fig.7a**). Immunohistochemical analysis confirmed a marked reduction in OTP protein signal in sgOtp - treated mice compared to sgCtrl controls (**Fig. 4b,c**).

**Fig. 4:**
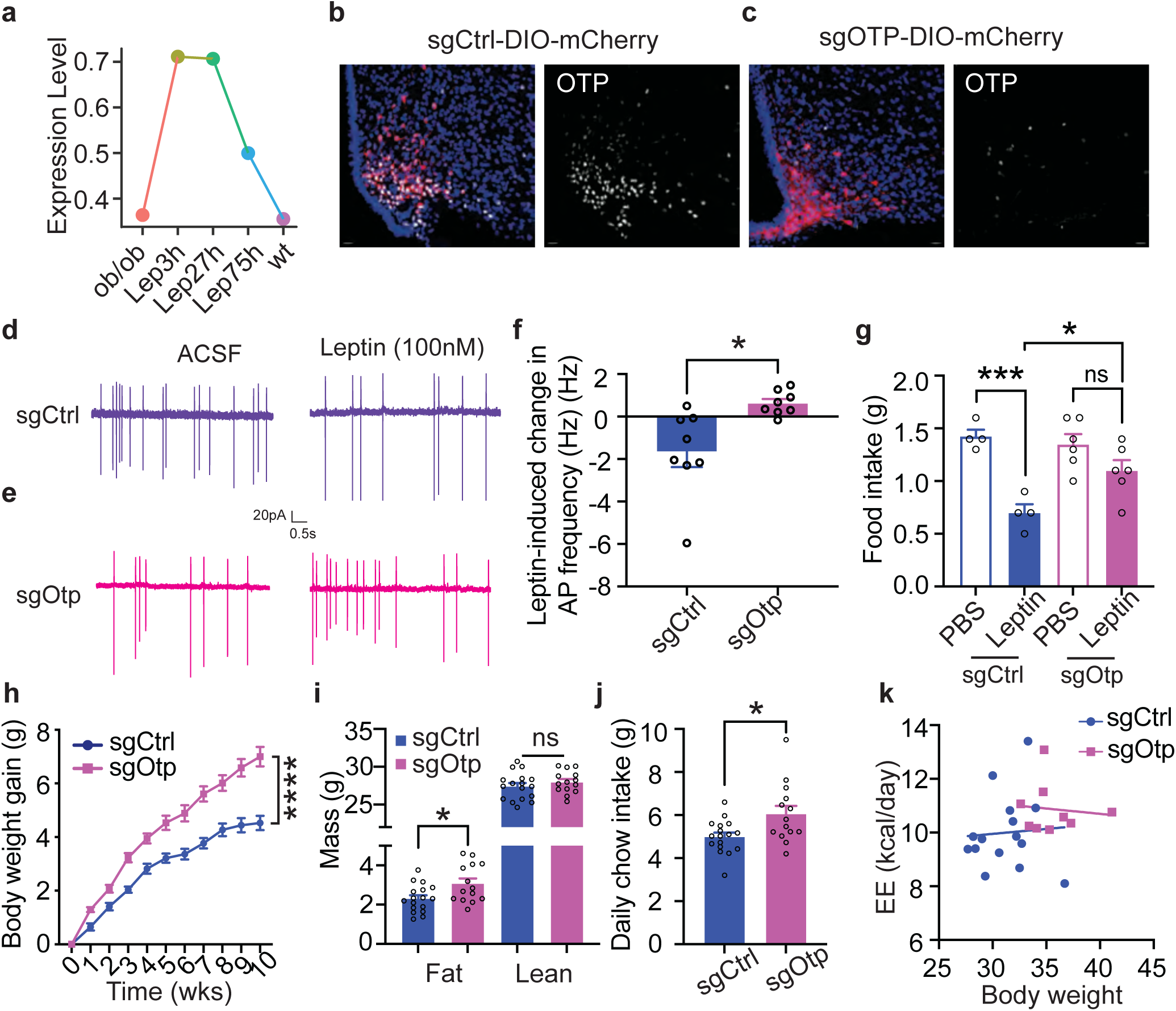
OTP is required for leptin-mediated inhibition of AGRP neurons. **a,** Otp expression in AGRP/LepR neurons across experimental conditions. **b,c,** Representative images of OTP immunostaining in AGRP neurons expressing sgCtrl (**b**) or sgOtp (**c**). mCherry labels virally transduced AGRP neurons. **d,e,** Representative cell-attached recordings from sgCtrl (**d**) and sgOtp (**e**) AGRP neurons before and after leptin application. **f,** Change in action potential firing frequency following leptin treatment in sgCtrl and sgOtp AGRP neurons. **g,** Food intake following PBS or leptin administration in sgCtrl and sgOtp mice. **h,** Body weight gain over time in sgCtrl and sgOtp mice. **i,** Lean and fat mass in sgCtrl and sgOtp mice at the study endpoint. **j,** Daily chow intake in sgCtrl and sgOtp mice. **k,** Energy expenditure (EE) plotted as a function of body weight in sgCtrl and sgOtp mice. Data are mean ± s.e.m. ns, not significant; *p<0.05; **p<0.01; ***p<0.001.

We first examined the acute electrophysiological response of AGRP neurons with an OTP knockout. In control neurons (sgCtrl), leptin robustly suppressed firing rates, consistent with canonical leptin-mediated inhibition. In contrast, AGRP neurons lacking OTP failed to reduce firing in response to leptin (**Fig. 4d-f**), demonstrating that OTP is required for leptin-induced suppression of AGRP neuronal activity. We next assessed the behavioral consequences of OTP loss in AGRP neurons. Following recovery from viral injection (2-3 weeks), mice received leptin or vehicle prior to the dark phase, and food intake was subsequently measured. Leptin significantly suppressed feeding in sgCtrl mice, but this acute anorectic effect was absent in sgOtp mice (**Fig. 4g**), indicating that OTP is necessary for leptin-induced satiety. Finally, we examined the long-term metabolic consequences of *Otp* disruption in AGRP neurons. Over a 10-week period following sgRNA injection, sgOtp mice gained significantly more weight than controls, with the increase in body weight consisting primarily of increased fat mass and no detectable change in lean mass (**Fig. 4h,i**). This weight gain was accompanied by elevated daily food intake (**Fig. 4j**). Although unadjusted energy expenditure (EE) was higher in sgOtp mice, ANCOVA analysis revealed no significant difference between groups after accounting for body weight (**Fig. 4k, Extended Data Fig. 7b**). Respiratory exchange ratio (RER) and locomotor activity were unchanged across the 24-h period and during both light and dark phases (**Extended Data Fig. 7c,d**), suggesting that increased food intake was the primary driver of the metabolic phenotype. Consistent with the increased adiposity, sgOtp mice also exhibited impaired glucose tolerance and insulin sensitivity at later time points following *Otp* disruption (**Extended Data Fig. 7e-h**). Together, these findings establish OTP as a cell-type-specific transcriptional regulator required for leptin-mediated inhibition of AGRP neurons and for regulating acute and chronic food intake and body weight.

### NOVA1 bidirectionally regulates energy balance in PVH neurons

We next applied a similar approach to identify functionally important leptin-mediated regulatory programs in downstream PVH neurons. Because PVH neurons predominantly function as a convergent satiety-promoting output node, and do not contain LepR-expressing neurons^16^, we focused on identifying region-level regulators associated with leptin-responsive neuronal state transitions. RNA velocity analysis across PVH neurons identified *Nova1* as one of the top velocity-associated genes linked to leptin-responsive state transitions (**Fig. 5a**). *Nova1* expression was elevated in PVH neurons in leptin-deficient *ob/ob* mice and was decreased following leptin treatment (**Fig. 5a**). Conversely, NOVA1 protein levels increased in the PVH following overnight fasting or chemogenetic activation of AGRP/LepR neurons (**Fig. 5b,c**), showing that it is downstream of ARC neurons and further suggesting that NOVA1 expression is regulated by an upstream orexigenic drive.

**Fig. 5:**
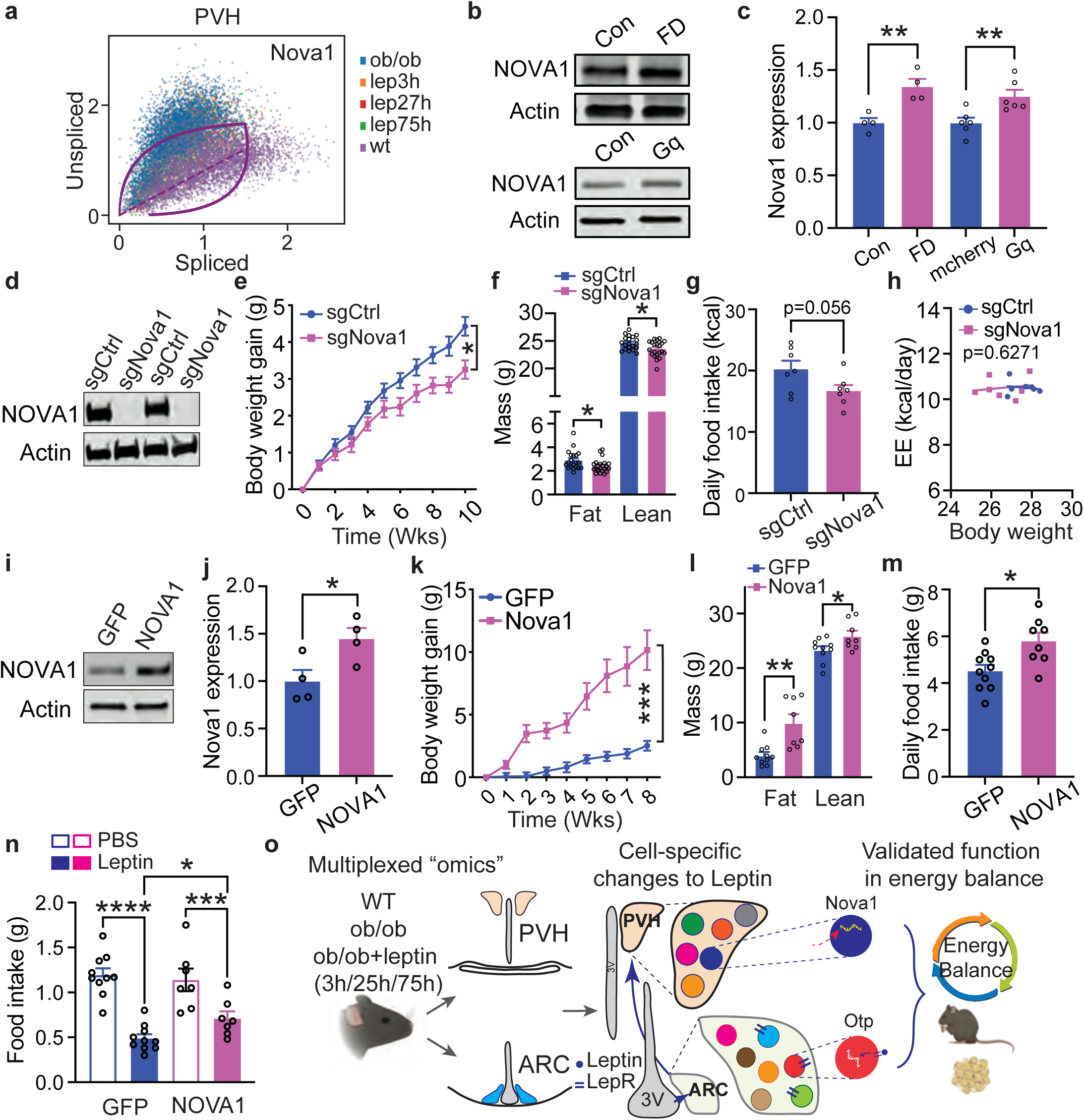
NOVA1 bidirectionally regulates energy balance in PVH neurons. **a,** Phase portrait of Nova1 in PVH neurons across experimental conditions. **b,c,** Representative western blots (**b**) and quantification (**c**) of NOVA1 protein levels in the PVH following overnight fasting (FD) or chemogenetic activation of AGRP neurons (Gq). **d,** Representative western blot showing NOVA1 protein levels in PVH tissue from sgCtrl and sgNova1 mice. **e,** Body weight gain in sgCtrl and sgNova1 mice. **f,** Lean and fat mass in sgCtrl and sgNova1 mice at the study endpoint. **g,** Daily food intake in sgCtrl and sgNova1 mice. **h,** EE plotted as a function of body weight in sgCtrl and sgNova1 mice. **i,j,** Representative western blot (**i**) and quantification (**j**) of NOVA1 protein levels in PVH tissue following AAV-mediated Nova1 overexpression. **k,** Body weight gain in GFP and NOVA1-overexpressing mice. **l,** Lean and fat mass in GFP and NOVA1-overexpressing mice at the study endpoint. **m,** Daily food intake in GFP and NOVA1-overexpressing mice. **n,** Food intake following PBS or leptin administration in GFP and NOVA1-overexpressing mice. **o,** Schematic summary of leptin-induced molecular state transitions across ARC and PVH neurons and functional validation of OTP and NOVA1 as regulators of energy balance. Data are mean ± s.e.m. *p<0.05; **p<0.01; ****p<0.0001.

To test this, we altered the level of NOVA1 expression in SIM1-expressing PVH neurons of otherwise WT mice. First, we selectively disrupted *Nova1* using a CRISPR-Cas9 strategy in which SIM1-Cre mice were mated to LSL-Cas9-GFP mice, and an AAV encoding sgRNAs targeting *Nova1* (sgNova1) or control sgRNAs was injected into the PVH. Western blot analysis confirmed a marked reduction in NOVA1 protein in sgNova1-injected PVH tissue compared to sgCtrl controls (**Fig. 5d**). Strikingly, mice lacking NOVA1 in PVH neurons gained significantly less body weight over a 10-week period, with reductions in both fat and lean mass (**Fig. 5e,f**). This reduced weight gain was accompanied by lower daily food intake (**Fig. 5g**), whereas ANCOVA analysis revealed no difference in energy expenditure between groups after accounting for body weight ( **Fig. 5h**). Consistent with this, EE, RER, and locomotor activity were unchanged across the 24-h period and during both light and dark phases (**Extended Data Fig. 8a-c**). Consistent with these findings, constitutive deletion of *Nova1* in SIM1-expressing cells using Sim1-Cre::Nova1^fl/fl^ mice also resulted in reduced body weight and food intake (**Extended Data Fig. 8d-f**).

To determine whether NOVA1 is sufficient to modulate feeding behavior, we next overexpressed NOVA1 in PVH SIM1 neurons using AAV-DIO-Nova1-GFP (**Fig.5i,j**). NOVA1 overexpression produced a robust increase in food intake and progressive weight gain, affecting both fat and lean mass (**Fig. 5k-m**). Moreover, NOVA1 overexpression attenuated leptin-induced suppression of food intake, indicating that elevated NOVA1 levels can blunt leptin’s satiety-promoting effects (**Fig. 5n**). Together, these findings show that NOVA1 levels in PVH are regulated by leptin and AGRP neuronal activity and that this protein bidirectionally controls feeding behavior and body weight.

## Discussion

Leptin is known to act on discrete populations of hypothalamic neurons to regulate feeding and other physiologic functions. However, our understanding of the cellular and molecular mechanisms through which leptin shapes hypothalamic feeding circuits is incomplete in part due to the extraordinary cellular diversity and functional complexity of its target regions. Here, we combined time-resolved snRNA-seq and snATAC-seq to systematically map leptin-induced changes in gene expression and chromatin accessibility over time across individual neurons in two key hypothalamic nuclei, the ARC, a direct leptin target, and the PVH, a secondary node. This integrated multi-omic framework revealed that leptin induces coordinated yet highly cell-type-specific transcriptional and regulatory state transitions that differ across neuronal populations and hypothalamic regions. Together, these findings provide a cell-resolved framework for understanding how a single hormonal signal engages distinct molecular programs across feeding circuits to control energy balance. The data also led to the identification of two genes, *Otp* and *Nova1*, that play a role in the function of these circuits.

Leptin induced widespread transcriptional and regulatory state changes in ARC and PVH without altering neural identity. Thus, the same cluster structure was maintained across a wide range of leptin concentrations, including in leptin-deficient, leptin-treated, and WT mice, indicating that leptin is not required for establishing or maintaining mature hypothalamic neuronal phenotypes. This observation is consistent with prior developmental studies showing that hypothalamic neuronal identities are specified during embryonic development, whereas leptin acts at later stages of circuit maturation to regulate their function^27–29^. Thus, leptin treatment induced substantial remodeling of intercellular communication, with distinct effects in LepR-expressing and non-LepR neuronal populations. In LepR-expressing neurons, leptin increased inferred interactions, consistent with enhanced network engagement. By contrast, non-LepR ARC neurons and PVH neurons exhibited reduced interaction numbers accompanied by more selective communication patterns, suggesting pruning of widespread low-specificity interactions is present in the leptin-deficient state. Together, these findings suggest that leptin promotes coordinated network refinement across feeding circuits rather than uniformly increasing neuronal connectivity.

In parallel, leptin induced substantial shifts in neuronal functional states, with the magnitude and coherence of these changes varying across regions and cell types. Leptin-induced state transitions were generally more pronounced in PVH neurons than in ARC neurons, consistent with a hierarchical organization in which leptin first engages ARC neurons and subsequently drives broader transcriptional remodeling in downstream PVH populations, which can be considered an effector site^2,10^. Although leptin treatment partially reversed transcriptional alterations associated with leptin deficiency, this rescue was incomplete. Most leptin-responsive populations showed an intermediate state between *ob/ob* and WT conditions, with relatively modest differences among the three leptin treatment durations examined. Even after 75 hours of leptin exposure, full convergence to the WT state was not observed, raising the possibility that longer and more chronic leptin exposure may be required to fully re-establish a WT state^30,31^. In addition, two ARC populations, clusters 16 and 18, marked by *Sim1* and *Sema3e*, respectively (**Fig. 3a**), showed little evidence of ordered leptin-responsive transitions, suggesting that certain neuronal types are either less sensitive to leptin and are likely to operate largely independently of leptin signaling^32^.

A longstanding question in leptin biology is how a single hormone acting through a shared receptor can elicit opposing physiological responses in distinct LepR-expressing neurons, such as hunger-promoting AGRP/LepR neurons and satiety-promoting BNC2/LepR or POMC/LepR neurons^3^. Our integrated multi-omic analyses reveal that leptin signaling through STAT3 is not implemented as a uniform transcriptional program across LepR neurons, but instead diverges in a highly cell-type-specific manner. In AGRP/LepR neurons, STAT3 preferentially regulated genes such as *Otp* and *Acvr1c*, which were not expressed in BNC2/LepR neurons. Conversely, in BNC2/LepR neurons, STAT3 induced a different set of genes, including *Prex2* and *Kctd*8, that were minimally expressed in AGRP neurons. This divergence provides a molecular framework for understanding how a shared signaling pathway can generate opposing functional outcomes in interspersed neuronal populations. How STAT3 selects distinct target genes across LepR neuron subtypes remains an open question. At a mechanistic level, this divergence is likely driven by differences in chromatin accessibility and cofactor availability, while these regulatory landscapes are themselves likely established by developmental lineage and cell-type-specific differentiation programs^33,34^. Notably, although leptin’s effects on neuronal firing occur on a rapid timescale that is unlikely to depend directly on STAT3-driven transcription^4,35^, our finding that loss of the STAT3 target OTP abolishes leptin-mediated inhibition of AGRP/LepR neuronal activity indicates that transcriptional programs may establish permissive neuronal states that enable rapid electrophysiological responses. Together, these results suggest that STAT3-dependent gene programs do not acutely mediate leptin signaling, but instead preconfigure specific neuronal states that enable rapid, cell-type-specific electrophysiological responses to leptin. Importantly, although these regulatory programs were identified through analyses in leptin-deficient mice, the functional consequences of manipulating key downstream effectors were validated in WT animals, establishing their relevance to physiological leptin action in intact feeding circuits. These included OTP and NOVA1, which were selected for further study because they were consistently prioritized across multiple complementary analyses, including differential expression, RNA velocity, regulatory network inference, regulation of gene expression by fasting and leptin and, in the case of OTP, regulation by STAT3.

OTP is a homeobox transcription factor that plays an essential role in hypothalamic development^25,36,37^. Germline loss of *Otp* disrupts neuronal differentiation in multiple hypothalamic nuclei, including the PVH, ARC, and supraoptic (SON) nuclei^25^, and germline *Otp* mutations have been linked to early-onset obesity in both mice and humans^38^, though its function in adults was not evaluated. The integrated transcriptomic, chromatin accessibility, and regulatory network analyses reported here identified OTP as a leptin-responsive regulator selectively induced by leptin in adult AGRP/LepR neurons. *Otp* emerged as a prominent velocity-associated gene, was predicted by SCENIC+ analysis to be regulated by STAT3, and was rapidly induced following leptin treatment. Functional studies further demonstrated that OTP is required for leptin-mediated inhibition of AGRP/LepR neuronal activity and feeding, as disruption of *Otp* in adult AGRP/LepR neurons abolished the electrophysiological response to leptin, blunted leptin-induced satiety, and promoted long-term weight gain. Together, these findings identify OTP as a critical mediator of leptin-responsive state transitions in hunger-promoting AGRP/LepR neurons and provide a potential mechanistic link between OTP dysfunction and obesity. Previous studies have demonstrated that OTP is required during development for the generation of AGRP neurons^26,33,39^, with AGRP⁺/NPY⁺ neurons largely absent in *Otp*-null mice or following developmental deletion of *Otp* in AGRP neurons. In contrast, in our study, adult deletion of *Otp* in AGRP neurons did not result in an obvious loss of AGRP-lineage cells over the experimental window examined. Using a Cre-dependent mCherry reporter in AGRP-Cre::LSL-Cas9 mice, we observed comparable numbers of labeled neurons in the ARC 10 weeks after sgOtp delivery (**Extended Data Fig 7a**), indicating that *Otp* is not required for the maintenance of AGRP-lineage neurons in adulthood. The use of scRNA seq to characterize gene expression in AGRP neurons with and without OTP expression can now be used to identify its target genes as well as reveal whether the expression of canonical AGRP identity markers (*Agrp*, *Npy*) is maintained in these cells and possibly the basis for the failure to respond to leptin in its absence. These and other studies will reveal how OTP contributes to maintaining a mature AGRP transcriptional state versus neuronal survival or other mechanisms.

The PVH is a known feeding center, though LepRb is not expressed there, as shown in previous studies and the data reported here^11,16^. It is thus a downstream node that is an important conduit for leptin signaling and provides an opportunity for further understanding how leptin’s signal is propagated. NOVA1 is a leptin-regulated gene expressed in PVH, and it was selected for further study because it was consistently identified across multiple complementary analyses, including differential expression, RNA velocity, and regulatory network inference. NOVA1 is an RNA-binding protein that regulates post-transcriptional gene expression, particularly alternative splicing, and has been extensively studied for its roles in neuronal development, synaptic function, and circuit balance, predominantly in inhibitory neurons^13,40^. Although recent work has linked NOVA1 to metabolic phenotypes^12^, its function within hypothalamic feeding circuits has remained largely unexplored. Through an unbiased, region-level analysis of leptin-responsive neuronal state transitions, we found that NOVA1 expression is dynamically modulated by energy state and leptin signaling. Importantly, manipulating NOVA1 expression in PVH SIM1 neurons bidirectionally altered food intake and body weight, establishing NOVA1 as a functional regulator of energy balance. Notably, NOVA1 is predominantly expressed in subcortical regions and has been studied primarily in *Gad2*⁺ inhibitory neurons given its strong enrichment in inhibitory neurons^41,42^, leading most prior work to focus on its role in inhibitory circuits. In contrast, our findings reveal an unexpected and essential role for NOVA1 in excitatory PVH SIM1 neurons, expanding the functional scope of NOVA1 beyond inhibitory neuron biology and suggesting a previously unrecognized post-transcriptional mechanism regulating hypothalamic control of feeding. As is also the case for OTP in the ARC, scRNA-seq of neurons after a PVH-specific knockout of NOVA1 may provide a basis for understanding how this protein influences the function of these neurons and feeding circuits.

Beyond OTP and NOVA1, our integrated analyses identified numerous additional candidate regulators across distinct leptin-responsive neuronal populations. For example, *Acvr1c* (activin A receptor type 1C), a predicted STAT3 target in AGRP/LepR neurons, has been genetically linked to obesity and is being explored as a therapeutic target for metabolic disease^43–45^. Likewise, *Alcam* (activated leukocyte cell adhesion molecule) emerged as a prominent leptin-responsive candidate in POMC/LepR neurons and has recently been implicated in hypothalamic regulation of energy balance^46–48^. In addition, *Ryr3* (ryanodine receptor 3) and *Lin28b* (lin-28 RNA-binding posttranscriptional regulator B), predicted STAT3 targets enriched in POMC/LepR neurons, have previously been implicated in metabolic regulation and obesity, respectively^49–51^. Together, these and other findings provide a rich resource for future mechanistic studies of leptin-responsive circuits. Further studies of these and potentially other genes in ARC and PVH, as well as leptin-responsive neuronal populations in other brain regions, may yield further information about how leptin signaling is propagated. In addition, because our analyses focused on congenital leptin deficiency with short-term leptin repletion, the extent to which the observed ARC-PVH state propagation and STAT3-divergent programs generalize to physiological or leptin-resistant obesity remains to be directly tested. Future studies applying similar multi-omic frameworks to diet-induced obesity or chronic leptin exposure will likely be informative for defining how these regulatory architectures adapt in leptin-resistant states.

In conclusion, our findings support a model in which leptin regulates energy balance not through a single uniform mechanism, but by engaging distinct layers of molecular control across hypothalamic circuits. By coupling cell-type-specific transcriptional regulation, chromatin remodeling, post-transcriptional control, and circuit-level communication, leptin establishes permissive neuronal states that enable rapid and context-dependent modulation of feeding behavior. This multi-scale regulatory architecture provides a framework for understanding how hormonal signals achieve both specificity and flexibility within complex neural circuits. More broadly, our study illustrates how integrated multi-omic approaches can reveal functionally meaningful regulatory nodes within heterogeneous brain regions and suggests new avenues for targeting discrete molecular programs to modulate energy balance.

## Methods

### Mice

All animal procedures were approved by the Institutional Animal Care and Use Committee at Rockefeller University and were conducted in accordance with the National Institutes of Health Guide for the Care and Use of Laboratory Animals. Mice were group housed under a 12-h light/12-h dark cycle at 22 °C and 30-60% relative humidity with ad libitum access to standard chow diet and water. The following mouse strains were used in this study: C57BL/6J (wild type; The Jackson Laboratory, stock no. 000664), Agrp-IRES-Cre (The Jackson Laboratory, stock no. 012899), Rosa26-LSL-Cas9 (The Jackson Laboratory, stock no. 024857), Lep^ob/ob^ (*ob/ob*, The Jackson Laboratory, stock no. 000632), Sim1-IRES-Cre (generated by Bradford Lowell, Beth Israel Deaconess Medical Center, Harvard Medical School), and Nova1^fl/fl^ mice (generated by Robert B. Darnell, Rockefeller University). Only heterozygous animals were used for all Cre-driver mouse lines. Sample sizes were selected based on previous studies using similar experimental approaches. Littermates of the same sex were randomly assigned to experimental or control groups when applicable. Recombinant mouse leptin (ThermoFisher Scientific, catalogue no. 498OB05M) was administered by intraperitoneal injection at 3 mg/kg to *ob/ob* mice and 5 mg/kg to non-*ob/ob* mice.

### Isolation of nuclei and multiplexed snRNA-seq and snATAC-seq

Male C57BL/6J and *ob/ob* mice (10-12 weeks of age) were euthanized by transcardial perfusion with ice-cold HEPES-sucrose cutting solution, as previously described. Brains were rapidly dissected, flash-frozen in liquid nitrogen, and stored at −80 °C until processing. Neuronal nuclei were isolated from frozen hypothalamic tissue and processed for multiplexed sn-RNA sequencing essentially as previously described. Briefly, nuclei were isolated by Dounce homogenization, stained with Hoechst 33342, an Alexa Fluor 647-conjugated anti-NeuN antibody, and TotalSeq hashtag antibodies, purified by fluorescence-activated cell sorting (FACS), and processed using the Chromium Single Cell 3′ Gene Expression v3 platform (10x Genomics). For multiplexed snATAC-seq, isolated nuclei were stained with 7-AAD, an Alexa Fluor 488-conjugated anti-NeuN antibody, and TotalSeq hashtag antibodies before FACS purification and library preparation using the Chromium Single Cell ATAC v2 platform (10x Genomics). Libraries were prepared by the Rockefeller University Genomics Resource Center and sequenced on Illumina or BGI platforms.

### Data analysis

#### snRNA-seq Data normalization, integration, and clustering

Raw FASTQ files were processed using Cell Ranger (v5.0, 10x Genomics) to generate gene expression and chromatin accessibility count matrices, respectively. Downstream analyses were performed using Seurat (v4.0.3)^52^, Scanpy^53^, and custom Python scripts. Cells with fewer than 800 or greater than 5,000 detected genes, greater than 12,000 total UMIs, or more than 1% mitochondrial transcripts were excluded. Genes detected in fewer than three cells were removed. Sample demultiplexing was performed using the Seurat hashtag oligonucleotide (HTO) workflow (positive quantile = 0.8), and only singlet nuclei were retained for downstream analyses. An additional low-quality cluster characterized by elevated mitochondrial transcript abundance and increased transcript counts, consistent with doublets and damaged nuclei, was identified and removed. Samples from different sequencing batches were integrated to correct batch effects via Canonical Correlation Analysis (CCA)^52^ before downstream analyses. Gene expression values were log-normalized (scale factor = 10,000), scaled, and subjected to principal component analysis (PCA). Uniform Manifold Approximation and Projection (UMAP) was used for dimensionality reduction and visualization. Cell clusters were identified using the Leiden algorithm with a resolution of 0.55. Seurat objects were subsequently converted to the h5ad format for downstream analyses in Scanpy. Cell clusters were annotated based on the expression of established marker genes and comparison with previously published hypothalamic single -cell transcriptomic atlases.

#### RNA velocity analysis

RNA velocity analysis was performed using scVelo^54^ to infer the directionality of transcriptional state transitions associated with leptin treatment. Spliced and unspliced transcript counts were obtained from loom files generated during read alignment. Expression matrices were preprocessed using the standard scVelo workflow, including gene filtering and normalization. A k-nearest-neighbor graph was constructed using the first 30 principal components and 30 nearest neighbors. RNA velocities were estimated using the stochastic model of transcriptional dynamics, and velocity graphs were computed to infer transition probabilities between nuclei. Velocity vectors were projected onto the precomputed UMAP embedding generated from the integrated transcriptomic analysis, enabling direct visualization of transcriptional dynamics within the same low-dimensional space used for clustering. Velocity fields were visualized to assess leptin-associated state transitions across neuronal populations.

#### Gene co-expression module analysis

Gene co-expression modules were identified using Hotspot^18^. Hotspot was applied to the full count matrix using the 5,000 previously identified highly variable genes. A depth-adjusted negative binomial model was used together with a weighted k-nearest-neighbor graph (k = 25) constructed from the PCA representation to estimate local gene autocorrelation. Genes with a false discovery rate > 0.05 were excluded from downstream analyses. Gene modules were identified by agglomerative clustering of the gene-gene similarity matrix. Module activity scores were calculated as the first principal component of each module and used as module eigenscores for downstream analyses.

#### Pseudo-bulk aggregation by batch and trajectory

For differential gene expression analyses, nuclei were aggregated into pseudo-bulk samples at the batch level within each major cell trajectory. Only batch–trajectory combinations containing at least 10 nuclei were retained. Gene-level counts were summed across nuclei to generate pseudo-bulk expression profiles. Counts were normalized using the trimmed mean of M-values method and transformed to log2 counts per million. Lowly expressed genes were filtered prior to downstream analyses.

#### Differential expression analysis across leptin conditions

Differential expression analysis was performed on pseudo-bulk expression profiles using the limma-voom framework. Mean-variance relationships were modeled using voom, and gene-wise linear models were fitted to assess transcriptional changes across leptin treatment conditions. Batch and other potential confounding variables were included in the design matrix when appropriate. Contrasts were specified to compare leptin-treated and control conditions within each major cell trajectory. Empirical Bayes moderation was applied to improve variance estimation. Genes with a false discovery rate (FDR) < 0.05 were considered differentially expressed.

#### Modeling leptin-associated gene expression trends

To characterize continuous transcriptional responses across leptin treatment conditions, generalized additive models were fitted to leptin-responsive differentially expressed genes. Models were fitted using an identity link function, a Gaussian error distribution, and cubic spline smoothing. Gene expression trends were evaluated across ordered leptin treatment conditions using evenly spaced prediction grids. Fitted expression profiles were normalized and hierarchically clustered using Ward’s method with Euclidean distance to identify groups of genes exhibiting similar leptin-response dynamics.

#### Eigentrend analysis

To summarize coordinated transcriptional dynamics of functionally related gene sets, eigentrends were computed from GAM-fitted expression trends. Fitted expression profiles were quantile-normalized across trajectories, and PCA was performed without centering. The first principal component, scaled between 0 and 1, was defined as the eigentrend. Statistical comparisons between eigentrends were performed using one-sided Wilcoxon signed-rank tests.

#### Gene Ontology enrichment analysis of gene trends

To interpret the biological processes underlying leptin-associated transcriptional dynamics, Gene Ontology (GO) enrichment analysis was performed on gene clusters identified from the GAM-based trend analysis. For each major trajectory, genes within each expression trend cluster were tested for enrichment of GO Biological Process and Molecular Function terms using a hypergeometric test with Benjamini-Hochberg FDR correction. The background gene set consisted of all genes expressed within the corresponding trajectory and retained after pseudo-bulk filtering. GO terms with an adjusted P < 0.05 were considered significantly enriched. Redundant GO terms were collapsed based on semantic similarity, and representative GO terms were used to define functionally coherent gene sets for downstream eigentrend analysis.

#### Cell-cell communication analysis

Cell-cell communication analysis was performed using CellChat^55^. A CellChat object was constructed from the normalized expression matrix and corresponding cell identity annotations, focusing on neuronal populations. Ligand-receptor interactions were inferred using the curated CellChat database, and communication probabilities were calculated using the CellChat statistical framework. Interactions supported by fewer than 10 cells per interacting group were excluded. Signaling networks were analyzed at both the ligand-receptor and signaling pathway levels to quantify interaction strength and communication patterns between sender and receiver populations. Condition-specific analyses were performed to compare leptin-treated and control groups, and signaling networks were aggregated and visualized using standard CellChat functions.

### snATAC-seq data preprocessing and quality control

#### snATAC-seq data processing and integration

Raw FASTQ files were processed using Cell Ranger ATAC (v2.1.0, 10x Genomics) against the mm10 reference genome to generate chromatin accessibility count matrices. Downstream analyses were performed using Seurat (v5.0.3) and Signac (v1.13.0). ATAC counts were normalized using term frequency-inverse document frequency (TF-IDF), and gene activity scores were calculated by aggregating chromatin accessibility over gene promoter regions. RNA and ATAC datasets were integrated using the nearest-neighbor label transfer workflow implemented in Seurat, enabling transfer of cluster identities from the RNA modality to the ATAC modality. The integrated datasets were co-embedded into a shared low-dimensional space for downstream analyses.

#### RNA-ATAC concordance analysis

To identify genes with concordant or discordant chromatin accessibility and transcriptional responses to leptin, per-condition average RNA expression (log-normalized counts) and ATAC gene activity scores were calculated for ARC AgRP/LepR neurons (cluster 0). Pearson correlation coefficients were computed between RNA and ATAC profiles across shared genes, and genes were ranked according to RNA-ATAC concordance. The top 500 most concordant and bottom 500 most discordant genes were visualized using row-scaled Z-score heatmaps, with hierarchical clustering performed on the RNA expression matrix and applied to both RNA and ATAC modalities.

#### SCENIC+ regulon analysis and STAT3 target gene identification

Transcription factor regulons were inferred using SCENIC+^24^, which integrates chromatin accessibility, gene expression, and transcription factor motif information to predict enhancer– gene regulatory relationships. STAT3 target genes identified by SCENIC+ were used for downstream expression and functional analyses. For visualization of STAT3 target gene expression, scaled RNA expression values were averaged by cluster and condition for ARC AgRP/LepR (cluster 0), POMC/LepR (cluster 3), and BNC2/LepR (cluster 17) neurons from wild-type mice. Genes with zero variance across cluster averages were excluded, and hierarchical clustering was performed to identify gene expression modules. GO over-representation analysis was performed for each module using compareCluster from clusterProfiler (v4.10.1) and the org.Mm.eg.db annotation database. Both Biological Process and Molecular Function ontologies were tested. To evaluate the robustness of enrichment results, analyses were repeated using three alternative background gene sets: the annotated mouse genome, Seurat highly variable genes, and all SCENIC+ peak-to-gene target genes. P values were adjusted using the Benjamini– Hochberg false discovery rate procedure.

#### STAT3 chromatin accessibility footprinting

STAT3 chromatin accessibility footprinting was performed using the Footprint function in Signac (v1.13.0). STAT3 binding motifs (JASPAR 2022, MA0144.1) were identified using TFBSTools (v1.40.0) and motifmatchr (v1.24.0). Hexamer insertion bias correction was applied using all peaks in the dataset (in.peaks = FALSE). Footprint analysis was performed on the merged ARC ATAC dataset, and Tn5 insertion profiles were visualized for AGRP/LepR (cluster 0), POMC/LepR (cluster 3), and BNC2/LepR (cluster 17) neurons individually and in combination. Tn5 insertion profiles were smoothed using a ±7 bp sliding window and visualized within ±100 bp of the motif center.

#### Analysis of regulons

snATAC-seq data from the ARC and PVH were processed using pycisTopic. Pseudo-bulk accessibility profiles were generated from Seurat-defined cell clusters, and consensus peaks were identified following MACS peak calling using the standard pycisTopic workflow^56^. Cells passing standard snATAC-seq quality control metrics were retained for downstream analyses. Topic modeling was performed using latent Dirichlet allocation, followed by Leiden clustering and UMAP visualization. Differentially accessible regions were identified using Wilcoxon rank-sum tests. Genome annotation and gene coordinates were obtained from the November 2020 archived Ensembl mouse genome (mm10).

Motif enrichment analysis was performed using pycisTarget with the Mus musculus (mm10) cisTarget databases. Chromatin accessibility, gene expression, and motif enrichment results were integrated using SCENIC+ to infer enhancer-driven regulons (eRegulons) describing transcription factor-regulatory element-target gene relationships. eRegulon activity scores were calculated and used to characterize regulatory programs across neuronal populations and leptin treatment conditions.

#### Viruses

AAV vectors used in this study were obtained from Addgene or generated through the Janelia Viral Tools Service. The following viral vectors were used: AAV5-EF1α-DIO-EYFP (Addgene #27056; 1.6 × 10^13^vg/mL). For *Otp* and *Nova1* deletion, sgRNA expression vectors were cloned in-house and packaged as AAV5 by the Janelia Viral Tools Service. The Otp sgRNA sequences were 5′-GTATCGGGCTGACCGAGTCG-3′ and 5′-GCCTGGCGCGCGGAACACGT-3′ (virus titer, 3.73 × 10^12^ vg/mL). The Nova1 sgRNA sequences were 5′-GATGCGATCAGGATTAACGG-3′ and 5′-ACCAAGAGGACCAACACGGG-3′ (virus titer, 2.47 × 10^12^ vg/mL). A non-targeting control sgRNA (5′-TTTTTTTTTTTTTTGAATTC-3′) was packaged as AAV5 (virus titer, 2.53 × 10^12^ vg/mL). For *Nova1* overexpression, an AAV5-hSyn-DIO-EGFP-P2A-Nova1 vector (virus titer, 3.88 × 10^12^ vg/mL) was cloned in-house and packaged by the Janelia Viral Tools Service. Viral aliquots were stored at −80 °C until stereotaxic injection.

#### Stereotactic surgery

Mice (8-10 weeks of age) were anesthetized with isoflurane (5% for induction and 1.5-2% for maintenance) and secured in a stereotaxic frame (Kopf Instruments, Model 1900). Viral vectors were delivered using a glass micropipette connected to a Nanoject III programmable nanoliter injector (Drummond Scientific). Bilateral injections (150 nL per site) were performed at a rate of 1 nL/s. Stereotaxic coordinates relative to bregma were as follows: arcuate nucleus (ARC), anteroposterior (AP), −1.65 to −1.70 mm; mediolateral (ML), ±0.25 to 0.30 mm; dorsoventral (DV), −5.9 mm; paraventricular hypothalamus (PVH), AP, −1.10 to −1.15 mm; ML, ±0.25 to 0.30 mm; DV, −4.9 mm.

#### Electrophysiology

Acute coronal hypothalamic brain slices (200 μm) were prepared from 2–3-month-old AGRP-Cre::LSL-Cas9-GFP mice 2–3 weeks after injection of sgCtrl or sgOtp virus into the ARC, using previously described procedures^57^. Mice were anesthetized with isoflurane before decapitation, and brains were rapidly removed and submerged in ice-cold slicing solution containing (in mM): 85 NaCl, 2.5 KCl, 0.5 CaCl₂, 4 MgCl₂, 25 NaHCO₃, 1.25 NaH₂PO₄, 64 sucrose, and 25 glucose. The solution was continuously bubbled with 95% O₂ and 5% CO₂ and maintained at pH 7.4. Coronal hypothalamic slices were prepared using a vibrating-blade microtome (VT1000S, Leica) and incubated at 32 °C for 40 min in artificial cerebrospinal fluid (ACSF) containing (in mM): 125 NaCl, 2.5 KCl, 1.25 NaH₂PO₄, 26 NaHCO₃, 10 glucose, 2 CaCl₂, and 1 MgCl₂, equilibrated with 95% O₂ and 5% CO₂ (pH 7.4). Slices were subsequently maintained at room temperature until recording.

Cell-attached patch-clamp recordings were performed at room temperature from mCherry-expressing neurons in the ARC. Neurons were identified by epifluorescence and visualized using differential interference contrast microscopy on an upright microscope (Axioskop 2FS Plus, Zeiss) equipped with a CCD camera (Hamamatsu). Patch pipettes were pulled from borosilicate glass (Sutter Instrument) using a PP-830 puller (Narishige), had resistances of 7–11 MΩ, and were filled with an internal solution containing (in mM): 135 potassium gluconate, 4 KCl, 0.05 EGTA, 10 HEPES, 4 MgATP, and 10 Na-phosphocreatine (pH 7.3, adjusted with KOH; 290 mOsm). Leptin was diluted in ACSF to a final concentration of 100 nM and bath-applied for 20 min by continuous perfusion following a baseline recording period. Recordings were acquired using an Axopatch 200B amplifier, low-pass filtered at 2 kHz, and digitized at 10 kHz using pClamp 10 software (Molecular Devices). Data were analyzed using IGOR Pro (WaveMetrics) and NeuroMatic. Firing rates were calculated in 10-s bins, and leptin-induced changes in firing rate were determined relative to the baseline firing rate for each neuron.

#### In situ hybridization

RNAscope fluorescent in situ hybridization was performed essentially as previously described. Briefly, mice were briefly transcardially perfused with ice-cold RNase-free PBS, and brains were rapidly collected, embedded in OCT, frozen on dry ice, and stored at −80 °C until sectioning. Coronal brain sections (15 μm) were collected onto Superfrost Plus Adhesion Slides (Thermo Fisher Scientific) and processed using the RNAscope Fluorescent Multiplex Assay (Advanced Cell Diagnostics) according to the manufacturer’s instructions. The following probes were used: Nos1 (cat. no. 437651-C2), Glp1r (cat. no. 418851-C2), Piezo2 (cat. no. 400191-C3), and Slc32a1 (cat. no. 319191). Sections were counterstained with DAPI and imaged using a Zeiss LSM 780 laser-scanning confocal microscope with a ×20 objective. Images were processed and analyzed using Fiji.

#### Immunohistochemistry and western blotting

Immunohistochemistry and western blotting were performed as previously described. For immunohistochemistry, mice were transcardially perfused with PBS followed by 4% paraformaldehyde (PFA). Brains were post-fixed overnight, cryoprotected in sucrose, embedded in OCT, and sectioned (30-50 μm). Free-floating sections were incubated with primary antibodies overnight at 4 °C, followed by fluorophore-conjugated secondary antibodies (Invitrogen). Sections were mounted onto Superfrost slides (Thermo Fisher Scientific) and imaged using a Zeiss LSM 780 laser-scanning confocal microscope with ×10 or ×20 objectives. Images were processed and analyzed using Fiji. The following primary antibody was used for immunohistochemistry: OTP (1:1,000; generated by Bora Lee and Jae W. Lee, University at Buffalo).

For western blotting, tissues were homogenized in lysis buffer supplemented with protease inhibitors. Equal amounts of protein were separated by SDS-PAGE, transferred onto PVDF membranes, and incubated with primary antibodies overnight at 4 °C, followed by fluorophore-conjugated secondary antibodies (LI-COR Biosciences). Protein bands were visualized using an LI-COR Odyssey Imaging System. The following primary antibodies were used: anti-NOVA1 (1:1,000; Abcam, ab183024) and anti-β-actin (1:2,000; Cell Signaling Technology, #3700S).

#### Indirect calorimetry

Indirect calorimetry was performed as previously described. Mice were individually housed in metabolic chambers (PhenoMaster, TSE Systems) with ad libitum access to food and water under a 12-h light/12-h dark cycle. Oxygen consumption, carbon dioxide production, food intake, locomotor activity, and energy expenditure were recorded continuously and analyzed using CalR.

#### Blood glucose, glucose tolerance, and insulin tolerance tests

Blood glucose measurements, glucose tolerance tests (GTT), and insulin tolerance tests (ITT) were performed as previously described. Blood glucose was measured using a OneTouch Ultra glucometer (LifeScan). For GTT, mice were fasted overnight and injected intraperitoneally with glucose (2 g/kg), and blood glucose levels were measured at 0, 15, 30, 60, and 120 min. For ITT, mice were fasted for 4 h before intraperitoneal injection of insulin (0.75 U/kg), and blood glucose levels were measured at 0, 15, 30, 45, and 60 min.

#### Statistical analysis

Statistical analyses were performed using GraphPad Prism 9 and R unless otherwise indicated. Data were tested for normality using the Shapiro-Wilk test, and appropriate parametric or non-parametric statistical tests were applied. All statistical tests were two-sided. Statistical methods used for each experiment are described in the corresponding Methods sections and figure legends. Biological replicates represent independent animals unless otherwise indicated. Detailed statistical results, including exact p-values, test statistics, degrees of freedom, and sample sizes, are provided in the Source Data. p < 0.05 was considered statistically significant unless otherwise indicated.

## Data availability

All data supporting the findings of this study are available within the manuscript, the Source Data, and the Supplementary Information. Raw and processed sn-RNA-seq and ATAC-sequencing data have been deposited in the Gene Expression Omnibus (GEO) under accession numbers GSE345448 (snRNA-seq) and GSE345945 (snATAC-seq).

## Code availability

Custom scripts used for data processing, analysis, and figure generation are available at https://github.com/RockefellerUniversity/Friedman_Han/tree/dev.

## Acknowledgement

We thank all members of the laboratory of J.M.F. for discussions and support. We also thank Professor Garry Nolan of the Department of Pathology at Stanford University for his support and mentorship of Y.T.

## Author contributions

H.L.T. conceived, designed, performed, and analyzed the research with input from all authors. H.L.T. and J.M.F. supervised the study. Y.T., J.R., and M.P. analyzed the mouse snRNA-seq and snATAC-seq data. Z.H., A.T., R.S., and A.I. contributed to the experiments. K.P. assisted with the metabolic cage experiments. L.K. performed the electrophysiology experiments and analyzed the data. B.L. provided the OTP antibody. Y.T. and R.B.D. provided the Nova1^fl/fl^ mouse line. P.C. provided guidance on metabolic phenotyping and input on the manuscript. H.L.T. and J.M.F. wrote the manuscript with input from all authors.

## Materials & Correspondence

Correspondence to Jeffrey M. Friedman or Han L. Tan.

---

**Supplementary Table 1: Ranked genes associated with RNA velocity in ARC neuronal populations.**

**Supplementary Table 2: Ranked genes associated with RNA velocity in PVH neuronal populations.**

**Supplementary Table 3: SCENIC+-predicted STAT3 target genes across LepR neuronal populations.**

**Extended Data Fig. 1:**
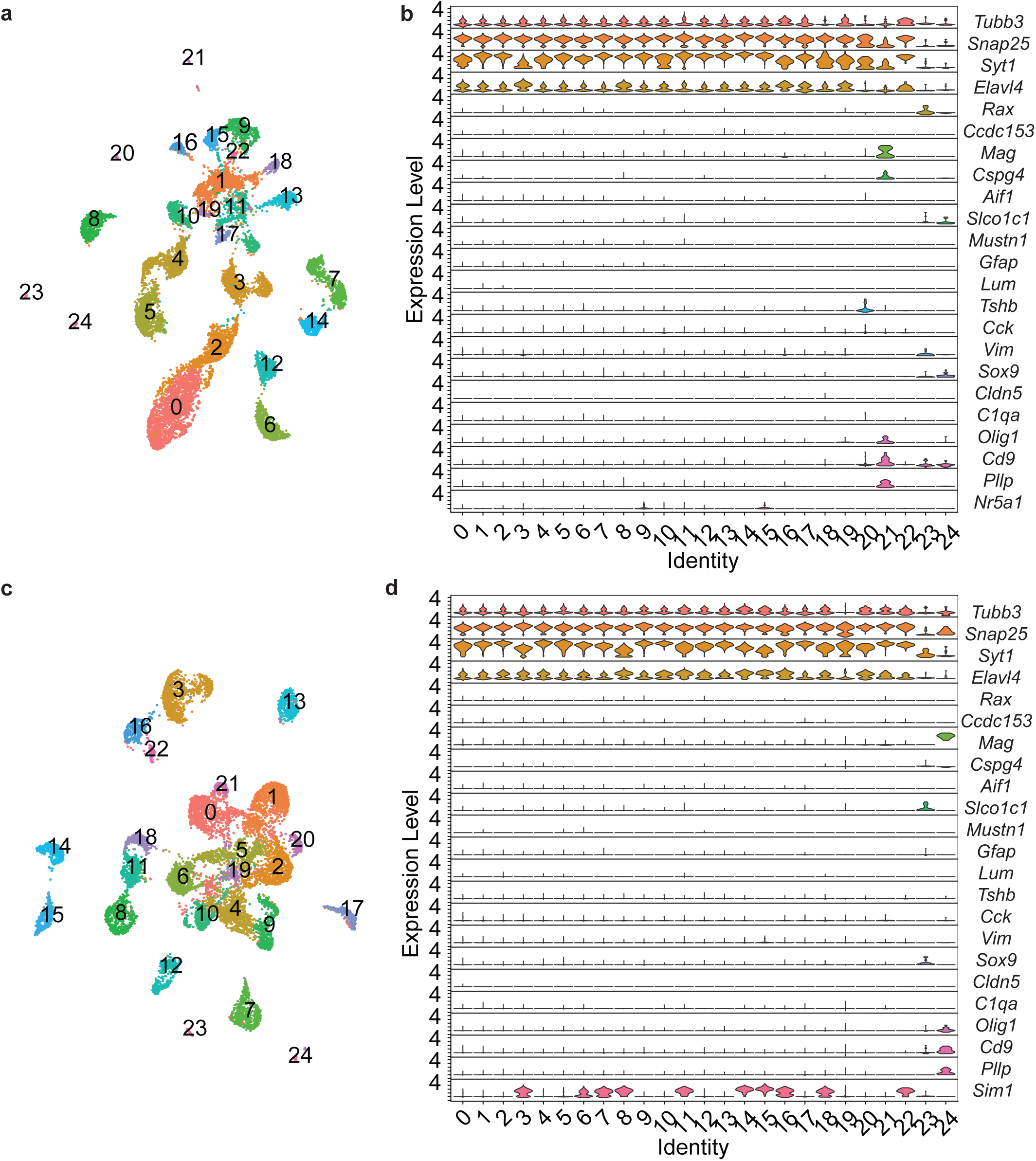
Cell-type annotation of ARC and PVH nuclei. **a,b,** UMAP visualization (**a**) and marker gene expression (**b**) of ARC cell populations identified by snRNA-seq. **c,d,** UMAP visualization (**c**) and marker gene expression (**d**) of PVH cell populations identified by snRNA-seq. Cell populations were annotated using established marker genes, including neuronal (*Tubb3*, *Snap25*, *Syt1*, and *Elavl4*), PVH neuronal (*Sim1*), tanycyte (*Rax* and *Cdol153*), ependymal (*Mag*), endothelial (*Cspg4*), astrocyte (*Aldh1l1* and *Slco1c1*), mural (*Mustn1*), fibroblast (*Lum* and *Dcn*), oligodendrocyte (*Olig1* and *Plp1*), and immune (*C1qa* and *Cd9*) markers.

**Extended Data Fig. 2:**
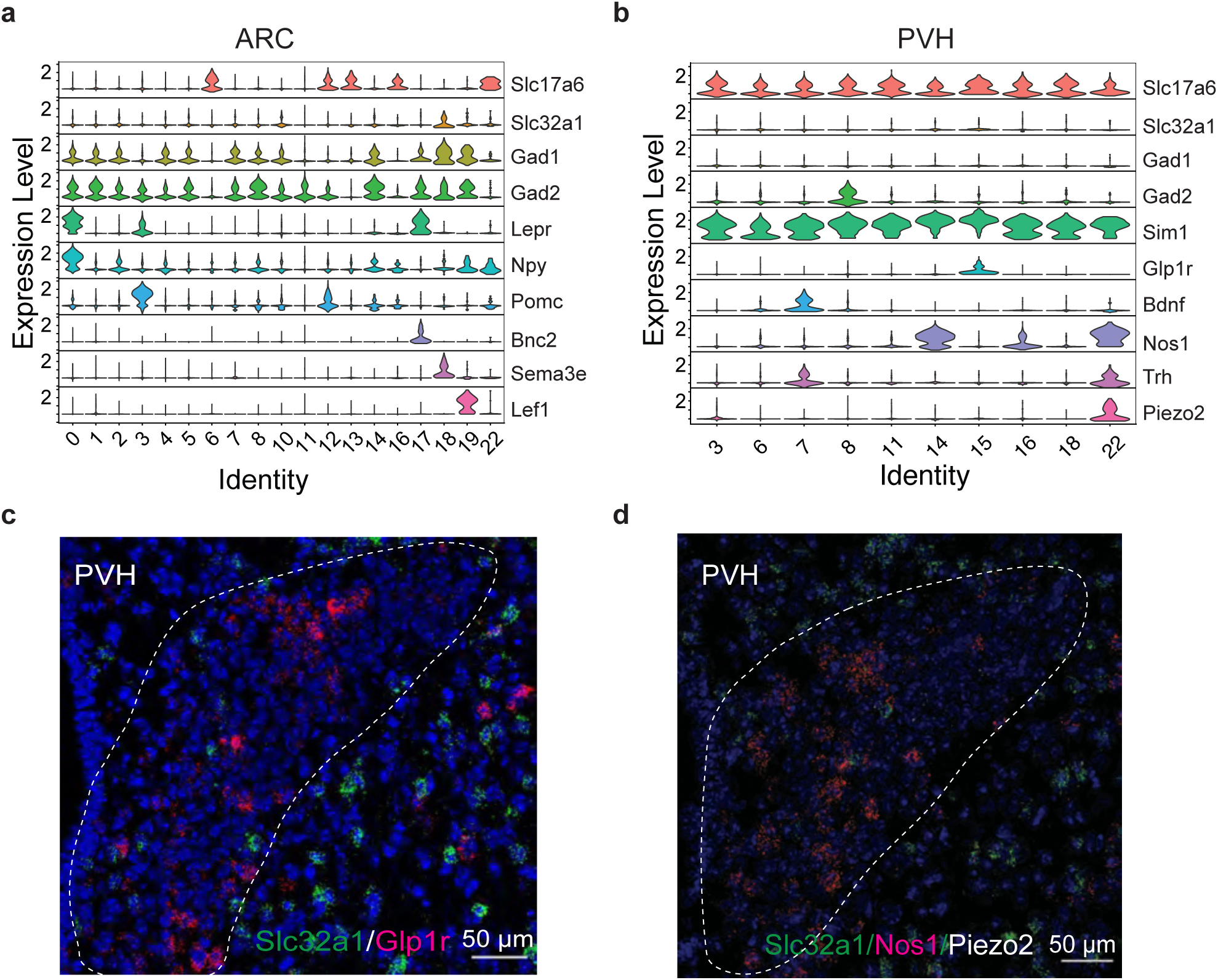
Molecular characterization of ARC and PVH neuronal populations. **a,b,** Expression of representative marker genes defining known neuronal populations across ARC (**a**) and PVH (**b**) neuronal clusters identified by snRNA-seq. **c,d,** Multiplexed RNAscope in situ hybridization validating Slc32a1-expressing and Piezo2-expressing cells in the PVH. Dashed lines indicate PVH boundaries. Scale bars, 50 μm.

**Extended Data Fig. 3:**
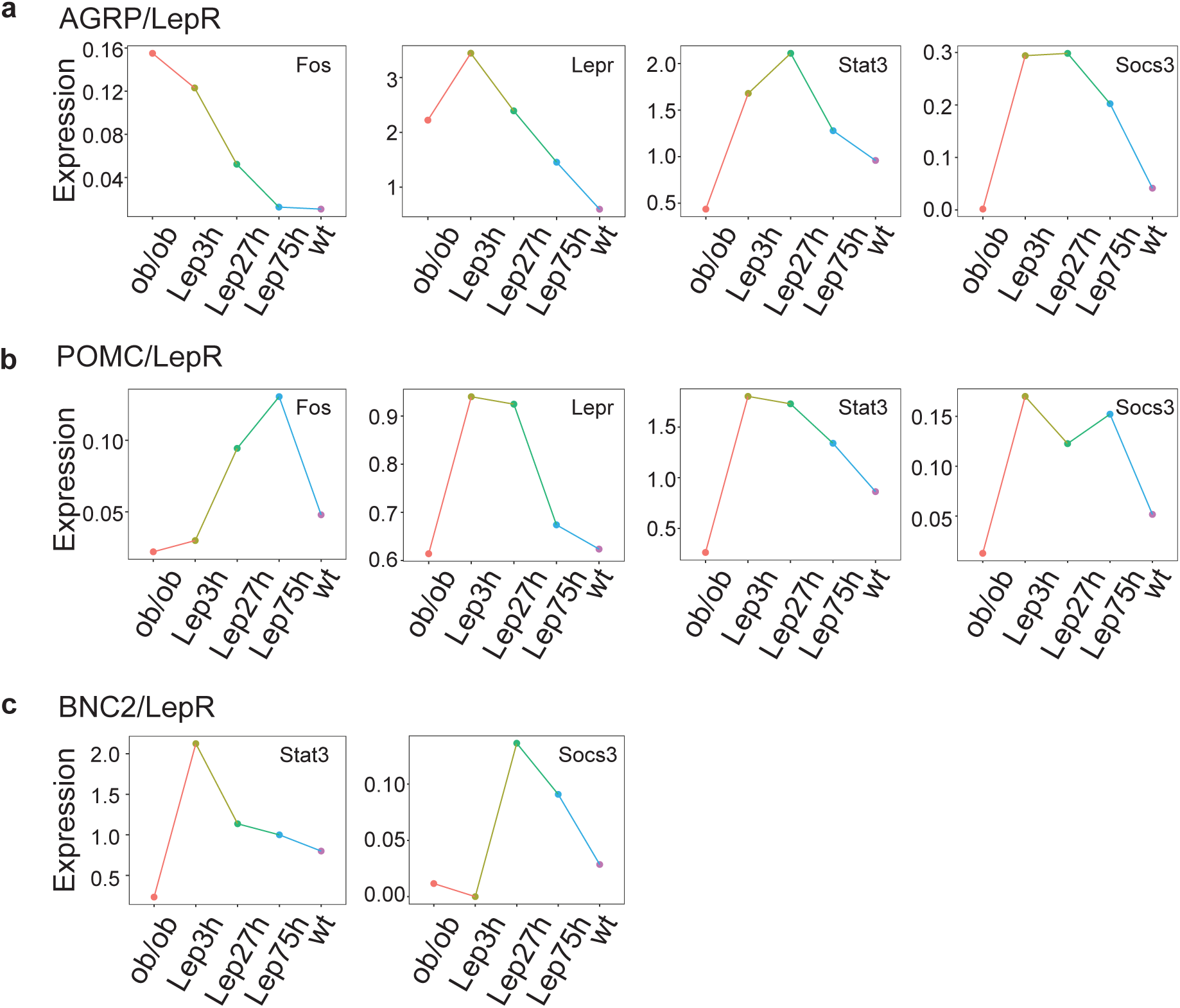
Expression dynamics of canonical leptin signaling genes. **a,b,c,** Expression of *Fos*, *Lepr*, *Stat3*, and *Socs3* across experimental conditions in AGRP/LepR (**a**), POMC/LepR (**b**) neurons, and BNC2/LepR (**c**) neurons.

**Extended Data Fig. 4:**
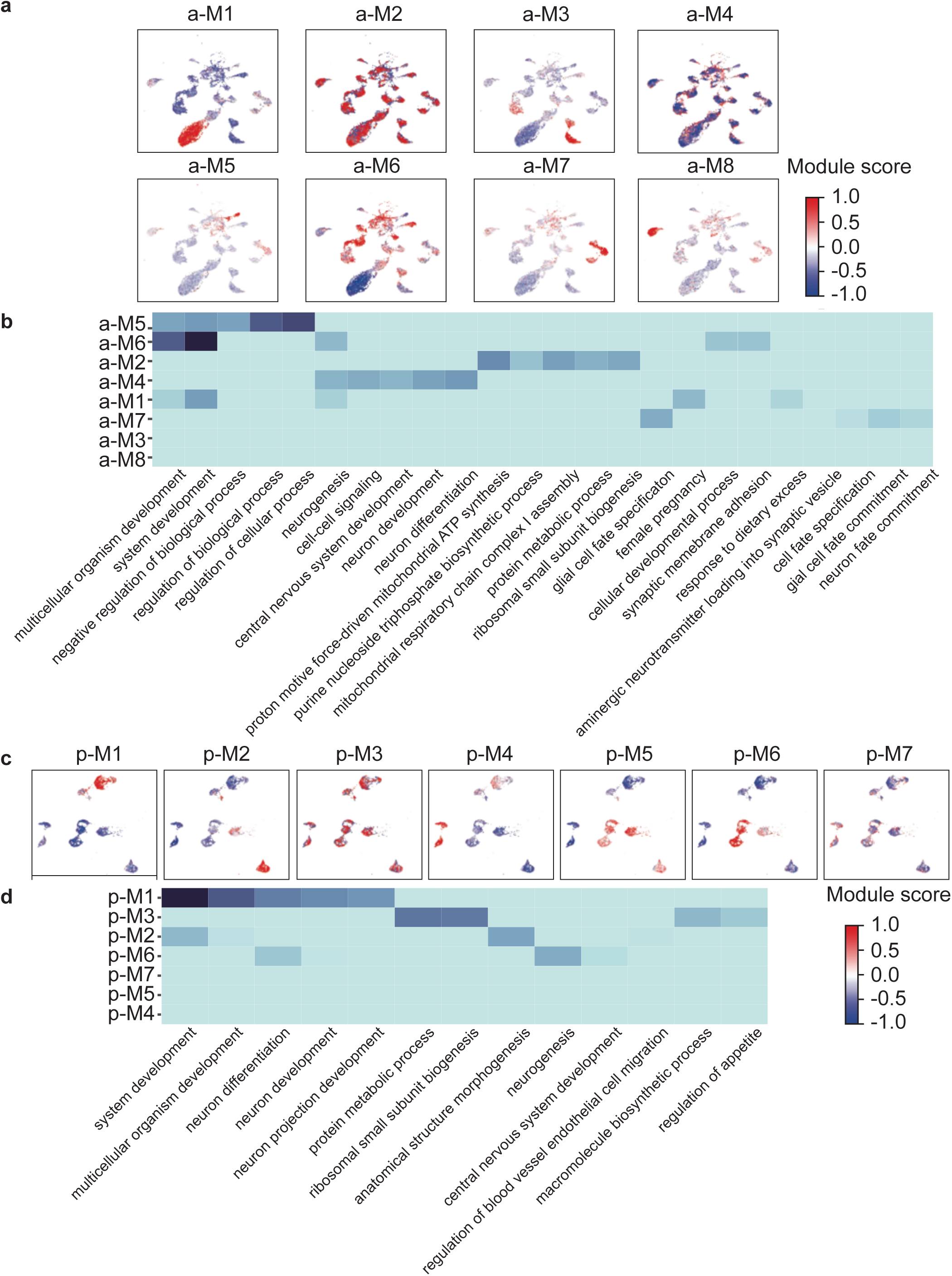
Hotspot analysis of ARC/PVH neuronal populations. **a,c,** Hotspot eigengene module scores overlaid on UMAP embeddings of ARC (**a**) and PVH (**c**) neurons. **b,d,** Top enriched Gene Ontology biological process terms associated with each module in ARC (**b**) and PVH (**d**).

**Extended Data Fig. 5:**
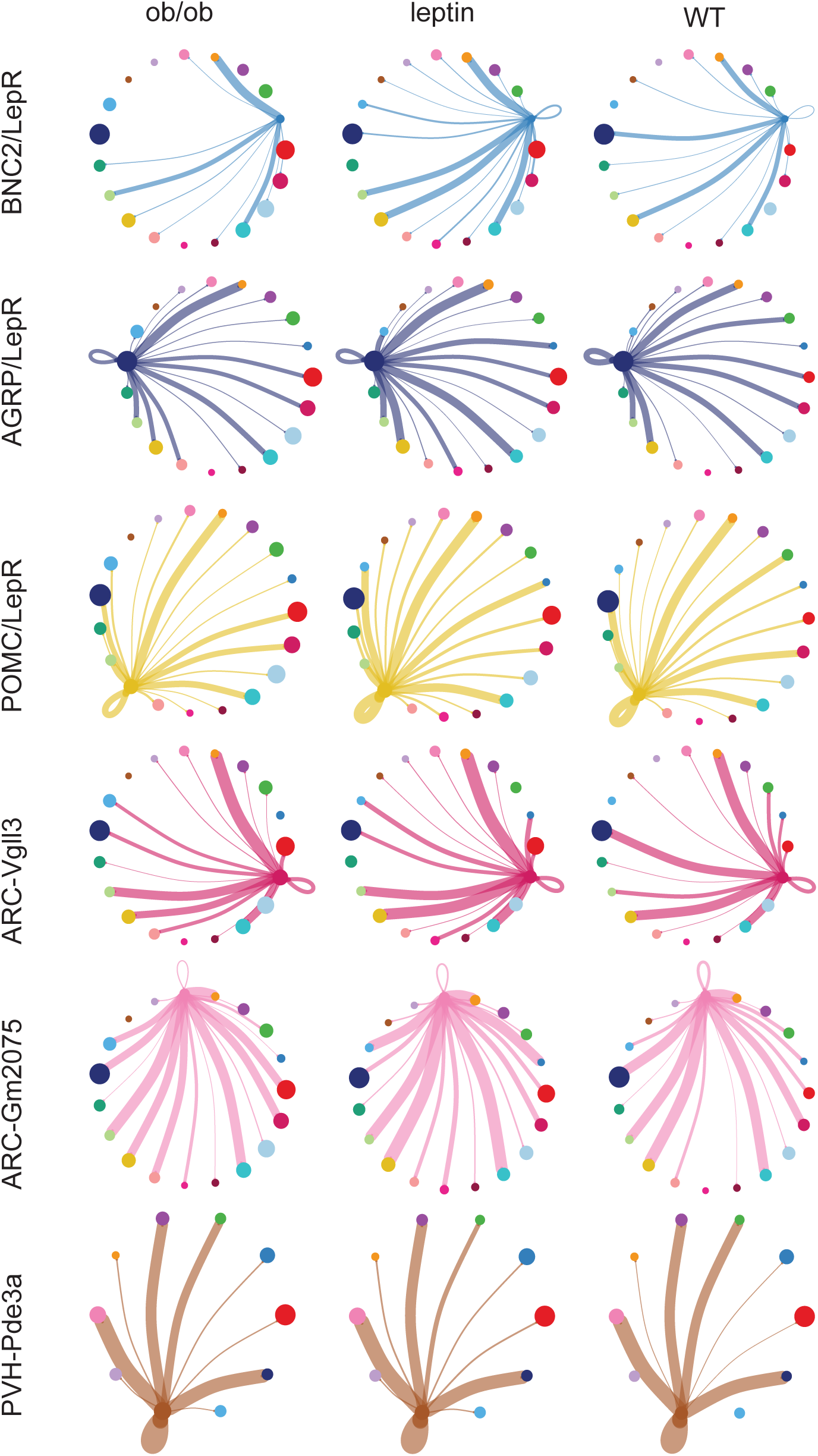
Cell-type-specific remodeling of intercellular communication following leptin treatment. Representative ARC LepR neurons, ARC non-LepR neurons, and PVH neuronal populations showing inferred cell–cell communication across ob/ob, leptin-treated ob/ob, and WT conditions. Each colored node represents an individual neuronal cluster, with colors corresponding to cluster identities across conditions. Connecting lines represent inferred cell–cell communication, with line thickness indicating communication strength.

**Extended Data Fig. 6:**
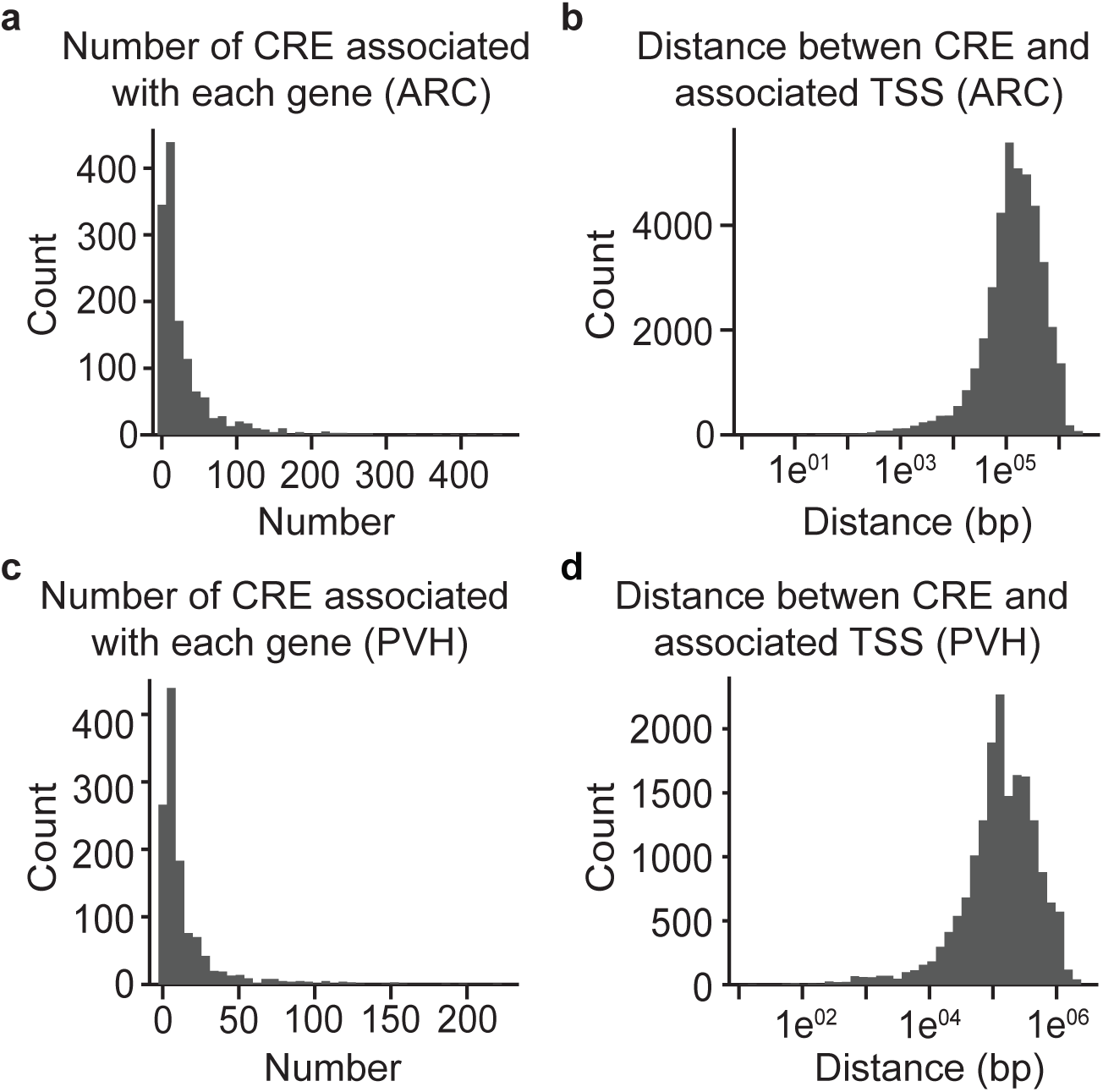
Characteristics of inferred cis-regulatory element–gene associations. **a,c,** Distribution of the number of cis-regulatory elements (CREs) linked to each gene in ARC (**a**) and PVH (**c**) neurons. **b,d,** Distribution of distances between linked CREs and the nearest transcription start site (TSS) of their associated genes in ARC (**b**) and PVH (**d**) neurons.

**Extended Data Fig. 7:**
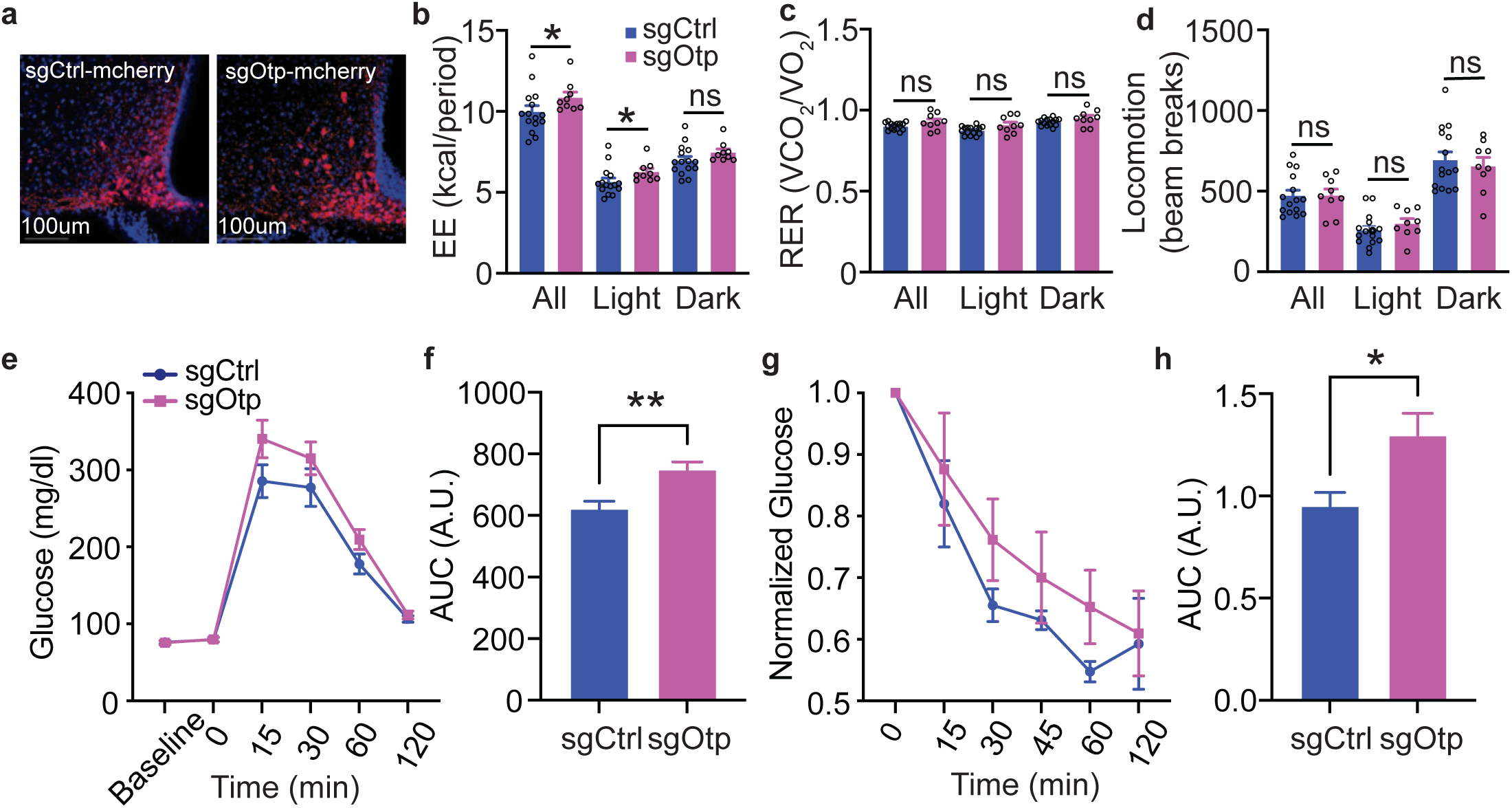
Metabolic characterization of mice with Otp deletion in AGRP neurons. **a,** Representative images of AGRP neurons expressing sgCtrl or sgOtp. mCherry labels virally transduced AGRP neurons. Scale bars, 100 μm. **b-d,** Energy expenditure (**b**), respiratory exchange ratio (RER; **c**), and locomotor activity (**d**) in sgCtrl and sgOtp mice, shown for the 24-h period (All) and separately for the light and dark phases. **e,f,** Glucose tolerance test (**e**) and quantification of the area under the curve (AUC; **f**) in sgCtrl and sgOtp mice at the study endpoint. **g,h,** Insulin tolerance test (**g**) and quantification of the area under the curve (AUC; **h**) in sgCtrl and sgOtp mice at the study endpoint. Data are mean ± s.e.m. *p<0.05; **p<0.01.

**Extended Data Fig. 8:**
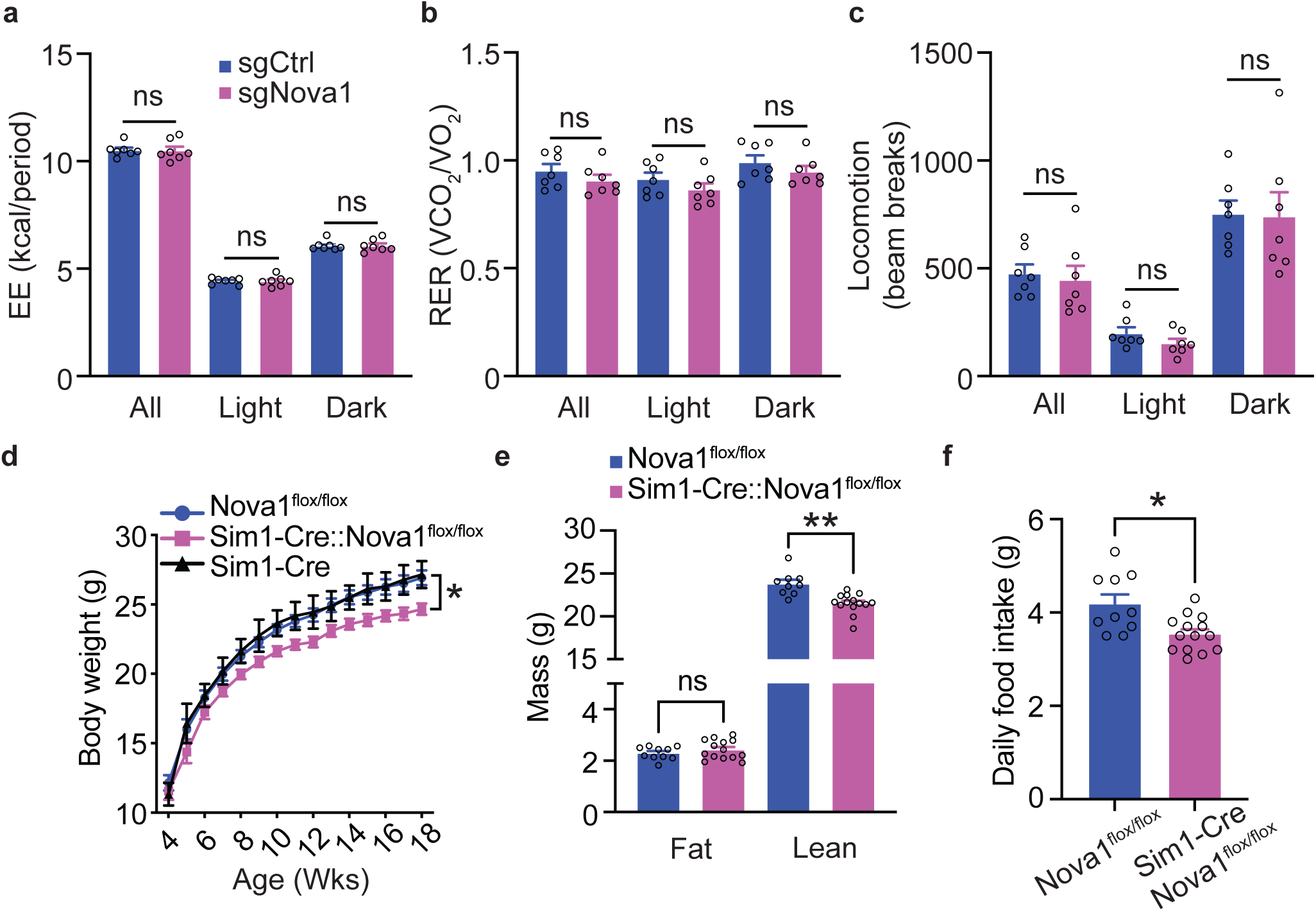
Deletion of Nova1 in SIM1 neurons reduces body weight and food intake. **a-c,** Energy expenditure (**a**), RER (**b**), and locomotor activity (**c**) in sgCtrl and sgNova1 mice, shown for the 24-h period (All) and separately for the light and dark phases. **d,** Body weight trajectories of Nova1flox/flox, Sim1-Cre, and Sim1-Cre::Nova1flox/flox mice. **e,** Lean and fat mass in Nova1flox/flox and Sim1-Cre::Nova1flox/flox mice at the study endpoint. **f,** Daily food intake in Nova1flox/flox and Sim1-Cre::Nova1flox/flox mice. Data are mean ± s.e.m. Data are mean ± s.e.m. *p<0.05; ***p<0.001.

